# A reproductive hierarchy controls mating fates among sibling yeast gametes

**DOI:** 10.64898/2026.10.17.759705

**Authors:** Chiara Vischioni, Benjamin P. Barré, Nina Vittorelli, Matteo De Chiara, Simone Pompei, Clément Vulin, Julien S. Luneau, Katarzyna Tomasiak, Nicolò Tellini, Simon Laugier, Sakshi Khaiwal, Sara Mitri, Bertrand Llorente, Marco Cosentino Lagomarsino, Gilles Fischer, Gianni Liti

## Abstract

Mating strategies shape inbreeding, gene flow and genetic diversity, yet their outcomes remain challenging to quantify. In *Saccharomyces*, population-genomic models predict mating predominantly between sibling meiotic products, with rare selfing and outcrossing. Here we developed an integrated framework combining multicolor flow cytometry, time-resolved imaging and mathematical modelling to directly quantify mating across scales. Bulk measurements revealed coexisting strategies whose frequencies vary with density, genotype and environment. A minimal stochastic model uses these population-level patterns to infer probabilistic hierarchies among mating modes, which single-tetrad analyses independently support. These reveal divergent and interdependent reproductive trajectories among sibling gametes, with one pair typically mating first and reshaping opportunities for the others. Temporal heterogeneity in germination and mating further diversifies reproductive outcomes and shapes compatibility, as experimental synchronization overcomes temporal reproductive barriers, enabling hybridization. Together, reproductive strategy emerges from heterogeneous gamete fates, raising the possibility that sibling diversification constitutes reproductive bet-hedging.

## Introduction

Despite its energetic cost, sexual reproduction is widespread across eukaryotes and profoundly shapes the long-term evolutionary success of a species, by generating the genetic diversity required for natural selection^1,2^. Yet, the driving forces under which organisms initiate sex, and the strategies they employ, remain among the most enduring puzzles in evolutionary biology^3^. Many organisms reproduce through alternative mating routes, including selfing, mating among relatives, and outcrossing, whose relative contribution can vary with the ecological and reproductive context. Such mating-system plasticity is well documented in plants^4,5^ and in self-fertile animals such as *C. elegans*^6,7^, where external cues can shift the balance between in- and out-breeding. Environmental control of reproductive strategy is particularly relevant to unicellular eukaryotes, which often inhabit unpredictable environments where resources are transient and information is limited. Many eukaryotic microbes engage in sexual reproduction only when the conditions become unfavorable, while proliferating clonally in nutrient-rich environments. Among them, *Saccharomyces* yeasts alternate between mitotic growth and sexual reproduction in response to nutrient availability (**Figure 1A**). Nitrogen and fermentable carbon starvation trigger meiosis, resulting in the formation of four haploid spores (gametes) enclosed in a protective ascus (the tetrad)^8^. Meiosis reshuffles the genome while halving its content. The haploid phase is typically transient: gametes restore their diploid state either by mating with a sibling spore within the same ascus (intra-tetrad mating), by fusing with an individual belonging to a distinct ascus (inter-tetrad mating, which constitutes outcrossing when partners are genetically distinct), or, if homothallic, by mating with their daughter upon mating-type switching (haplo-selfing)^9^. The route ultimately followed depends on the spatiotemporal context in which compatible partners are encountered^10,11^. Mating thus represents a decisive transition in the life cycle, functioning as a genomic reset that restores diploidy by determining which genomes are brought together in the resulting zygote, where genetic incompatibilities can compromise viability or fertility. Although the molecular mechanisms underlying yeast mating have been deeply characterized in laboratory settings^12^, our understanding of how mating strategies evolved and are shaped by environmental factors remains limited^13^. A major unresolved question is how the relative contribution of alternative reproductive routes varies across conditions. More fundamentally, it remains unclear whether population-level patterns reflect a reproductive strategy fixed at the level of a meiotic cohort or emerge from alternative fates realized by individual sibling gametes.

**Figure 1.**
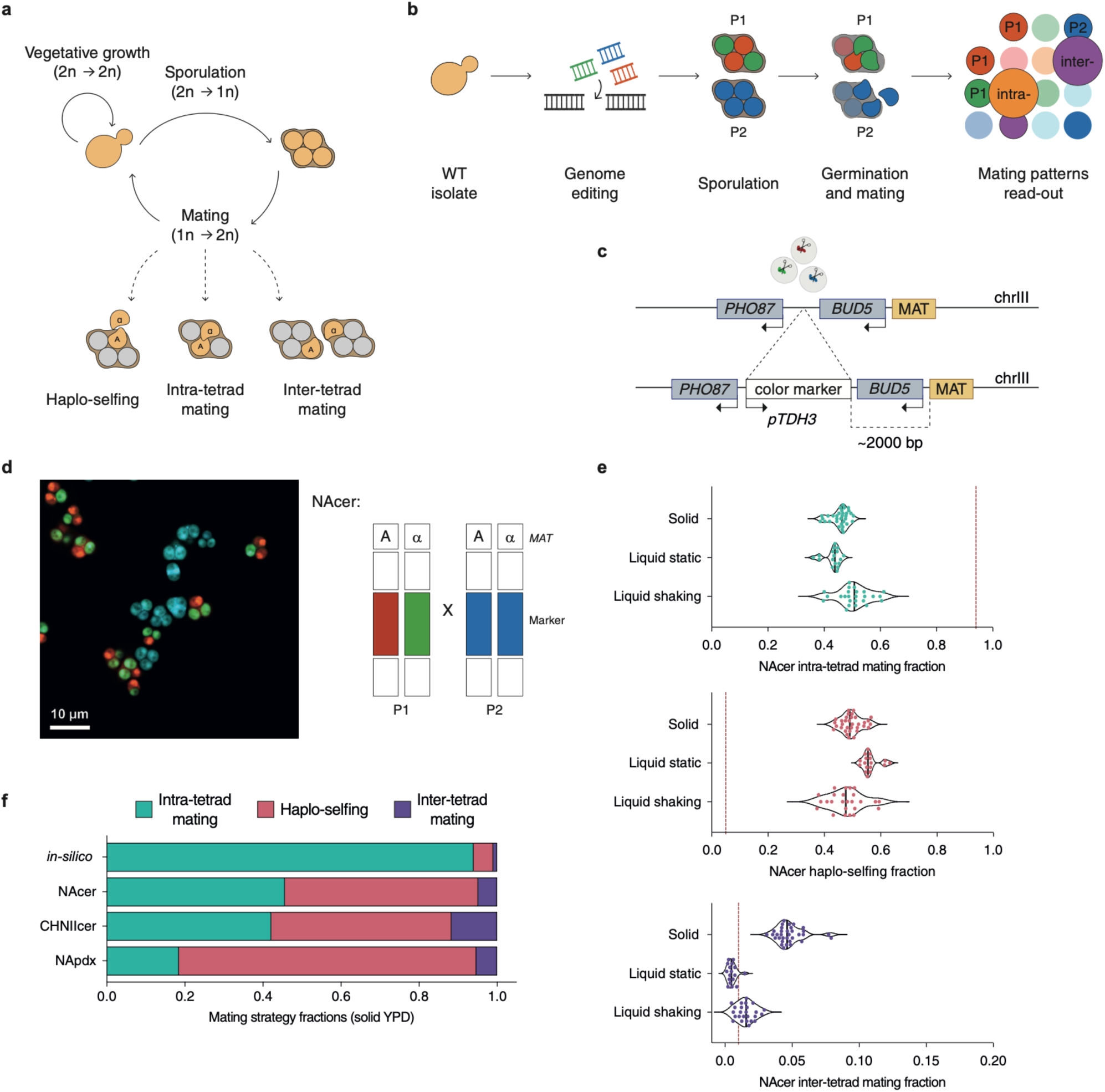
Direct measurement of yeast mating. **a,** Schematic representation of *Saccharomyces* life cycle highlighting transitions between vegetative proliferation (diploid → diploid), sporulation (diploid → haploid), and mating (haploid → diploid). Upon germination, spores restore diploidy through three possible routes: intra-tetrad mating between sibling spores within the same ascus, inter-tetrad mating between spores from unrelated asci, or haplo-selfing following mating-type switching in homothallic strains. **b,** gaMATe overview. Wild isolates were engineered to carry constitutively expressed fluorescent markers and then sporulated to generate tetrads. Mixed asci populations were allowed to germinate and mate under defined environmental conditions. Mating outcomes were quantified by flow cytometry based on fluorescent marker combinations, and mating frequencies were inferred using a mathematical framework (**Appendix 1**). **c-d,** CRISPR–Cas9 strategy for engineering the gaMATe system. Fluorescent markers were integrated using CRISPR/Cas9 at a neutral locus on chromosome III (designated as *rainbow* locus), linked to the mating-type locus (*MAT*). Each parental strain carries one or two constitutively expressed fluorescent reporters. e, Violin plots show the distribution of intra-tetrad mating, haplo-selfing and inter-tetrad mating fractions measured in NAcer under different physical conditions (solid agar, static liquid and shaken liquid YPD). Each point represents an independent measure (*N = 33* in solid agar, *N = 17* in liquid static, *N = 25* in liquid shaken). Central bars indicate the median. The vertical dashed line indicates mating frequency inferred from population genomic analyses^14^. Differences between mating distributions were quantified using Jensen–Shannon distance (JSdist; **Table S3; Figure S4**). **f,** Mating strategies vary across genetic backgrounds. Stacked bar plots showing the compositional distribution of mating outcomes across three *Saccharomyces* genetic backgrounds germinated and mated in solid YPD. Bars represent the mean mating fractions (individual replicates and SDs are shown in **Figure S2D**; *N = 33* in NAcer, *N = 16* in CHNIIcer, *N = 22* in NApdx). The top bar shows the compositional distribution of mating routes inferred from population genomic analyses^14^. Similarity in compositional patterns was quantified using Jensen-Shannon distance (JSdist = 0.11 for NAcer-CHNIIcer; JSdist = 0.25 for NAcer-NApdx; JSdist = 0.26 for CHNIIcer-NApdx; **Table S3; Figure S4**).

Estimates of the relative contributions of mating strategies have been primarily inferred from population genomic patterns rather than directly quantified. A seminal study on wild *S. paradoxus* populations estimated that approximately 94% of matings occur within tetrads, with haplo-selfing and outcrossing between unrelated lineages contributing only marginally^14^. These estimates, derived from genomic signatures of inbreeding, have profoundly influenced our understanding of yeast reproduction, and have contributed to a prevailing model of predominantly intra-tetrad mating across *Saccharomyces* species. However, large-scale population surveys have revealed unexpectedly high levels of heterozygosity and footprints of admixture across natural populations^15^, suggesting that outcrossing and hybridization may play a more substantial role than previously thought. Similarly, evidence of ancient introgression between *Saccharomyces* species indicates recurrent gene flow across lineages^16–18^. Together, these genomic patterns integrate mating history with subsequent evolutionary processes, shifting the question beyond which route dominates over evolutionary timescales towards when and under what conditions alternative reproductive outcomes are realized at ecological scale. Early experimental attempts to quantify yeast mating reported that 75–80% of spores mate within the same tetrad immediately after germination^19^, aligned with other population surveys of wild isolates inferring outcrossing to be rare^20^. The physical confinement within the ascus, reinforced by interspore bridges, is thought to enforce mating among meiotic siblings by maintaining spatial proximity^21^, thereby limiting opportunities for outcrossing. Yet ecological perturbations can dramatically increase outcrossing, for example through dispersal in insect guts^22,23^, or controlled inter-ascus contact^24^. These contrasting observations indicate that mating outcomes are highly context-dependent, highlighting the limitations of existing approaches for quantifying their relative contributions. Most estimates derive either from genomic signatures accumulated over long evolutionary timescales or from small-scale assays that cannot systematically disentangle intra-tetrad, haplo-selfing and inter-tetrad across diverse environments and genetic backgrounds, nor resolve the eventual fate of individual gametes.

Here, we address these gaps by developing a highly sensitive molecular system based on a multicolor flow cytometry design that, together with a validated mathematical framework, enables direct quantification of alternative yeast mating strategies under defined conditions. Across genetic and environmental contexts, we find that these routes coexist in shifting proportions rather than being dominated by a single reproductive outcome, while a minimal stochastic model reveals preferred probabilistic hierarchies among reproductive modes. Single-tetrad analyses further show that sibling gametes from the same meiosis frequently realize distinct reproductive fates. Together, our findings establish reproductive strategy as a plastic and emergent property of individual gamete behavior, raising the possibility that diversification among meiotic siblings provides a form of reproductive bet-hedging under uncertain mating opportunities.

## Results

### Direct quantification of yeast mating reveals a mixed spectrum of strategies

Population-genomics estimates have long described *Saccharomyces* sexual reproduction as dominated by intra-tetrad mating with the fusion of sibling gametes^14^. To experimentally test this assumption, we developed **gaMATe** (**ga**mete **MAt**ing **Te**chnology), a genetically engineered system based on a multicolor fluorescence design that enables a precise measure of all mating outcomes (**Figure 1B**, **Figure S1**). Our tool enables targeted CRISPR/Cas9 integration of one or two constitutively expressed fluorescent markers at a neutral site on chrIII (**Figure 1C-D**). This site lies ∼2kb to the left of the mating-type locus (MAT) (similar to^25^), minimizing recombination events that could unlink fluorescence from mating identity (0.5 - 1%, **Figure S2A**; **Methods**). Notably, gaMATe is broadly applicable, being readily compatible with 96% of the known *S. cerevisiae* genetic diversity (**Figure S1C**; **Methods**), and adaptable to all *Saccharomyces* species. We first applied gaMATe to our focal strain, the North American *S. cerevisiae* (YPS128, hereafter NAcer), a fully homozygous wild isolate typified by its high sporulation efficiency (>99% at 72 hours upon 2% potassium acetate induction)^15,26,27^. We developed and validated a mathematical framework to derive quantitative frequencies of mating events from observed fractions of cells carrying different colors (**Appendix 1, Figure S1**). The framework is based on a set of three equations relating the observed proportion of fluorescent populations to the frequencies of the different mating strategies (**Figure S1B**). These allow us to estimate the fraction of cells undergoing intra-tetrad mating, haplo-selfing, and inter-tetrad mating in the population, corrected for both the experimentally measured recombination rate and the same color combination profile arising from different mating outcomes (**Appendix 1, Figure S1A-B**).

Contrary to the canonical paradigm^14^, in NAcer, intra-tetrad mating is substantially less frequent than previously assumed (mean = 45.6% ± 3.1%, on solid YPD). In contrast, haplo-selfing was found at a higher-than-expected fraction (mean = 49.6% ± 3.6%, on solid YPD), and inter-tetrad mating, typically thought to be extremely rare, also contributed to a meaningful proportion of the mating pattern (mean = 4.8% ± 1%, on solid YPD) (**Figure 1E, Table S2-S3**). Together, these observations reveal a reproductive program comprising a combination of mating modes rather than one dominated by intra-tetrad fusion. To test how the physical environment influences mating outcomes, we measured NAcer mating in solid agar, static liquid, and shaken liquid YPD. While mating event frequencies differed, the mixed system was observed across all experimental set-ups (**Figure 1E, Table S2**). Static liquid cultures, where spores mostly remain clustered, but pheromone gradients are less maintained, favored haplo-selfing (mean = 56.3% ± 3%) and reduced both intra- and inter-tetrad mating (mean = 43.2% ± 2.9% and 0.5% ± 0.3% respectively) relative to solid agar. Shaken liquid promoted spore mixing and enhanced mating between non-sister gametes (mean = 1.5% ± 0.7%), whereas solid agar, where asci germinate in close spatial proximity and pheromone gradients are stable, yielded the highest inter-tetrad mating rate among the conditions tested, consistent with efficient local signaling and shmoo orientation^28,29^.

We next asked whether these patterns are conserved across wild lineages, applying gaMATe to two genetically distinct *Saccharomyces* backgrounds: a highly divergent *S. cerevisiae* isolate from China (SX3, hereafter CHNIIcer), and the undomesticated sister species *S. paradoxus* (the North American YPS138, hereafter NApdx)^26,30,31^. Generally, the environmental influence was consistent across backgrounds (**Figure S3**), whereas the quantitative balance among strategies varied markedly with the genetic background (**Figure 1F, Figure S2D, Table S2-S3**). At the same starting population density, in solid YPD, CHNIIcer showed higher inter-tetrad mating than NAcer (mean = 11.7% ± 2.6%), whereas between-strains differences in intra-tetrad mating and haplo-selfing were smaller (mean = 42.1% ± 3.3%, and 46.2% ± 3.4% respectively). By contrast, NApdx displayed a pronounced shift toward haplo-selfing (mean = 76.2% ± 8.5%) and reduced intra-tetrad mating proportion (mean = 18.5% ± 6.3%). These differences likely reflect background-specific variation in traits such as germination dynamics, coordination in gamete release, and spore viability, altering the encounter probability among sibling and non-sibling haploids. The mixed mating system uncovered here is a conserved yet quantitatively context-dependent feature, revealing an unexpectedly complex reproductive landscape in a genus long considered to be almost exclusively inbred via intra-tetrad mating.

### Rewiring yeast mating through genetics and ecological perturbations

The coexistence of multiple mating modes raises the question of whether their relative contributions can be reprogrammed by both genetic and environmental factors. To address this, we perturbed mating across successive levels of control corresponding to the constraints experienced by germinating spores: physical determinants of encounter geometry, signaling dynamics governing mating commitment, and genetic control of mating-type switching (**Figure 2A**).

**Figure 2.**
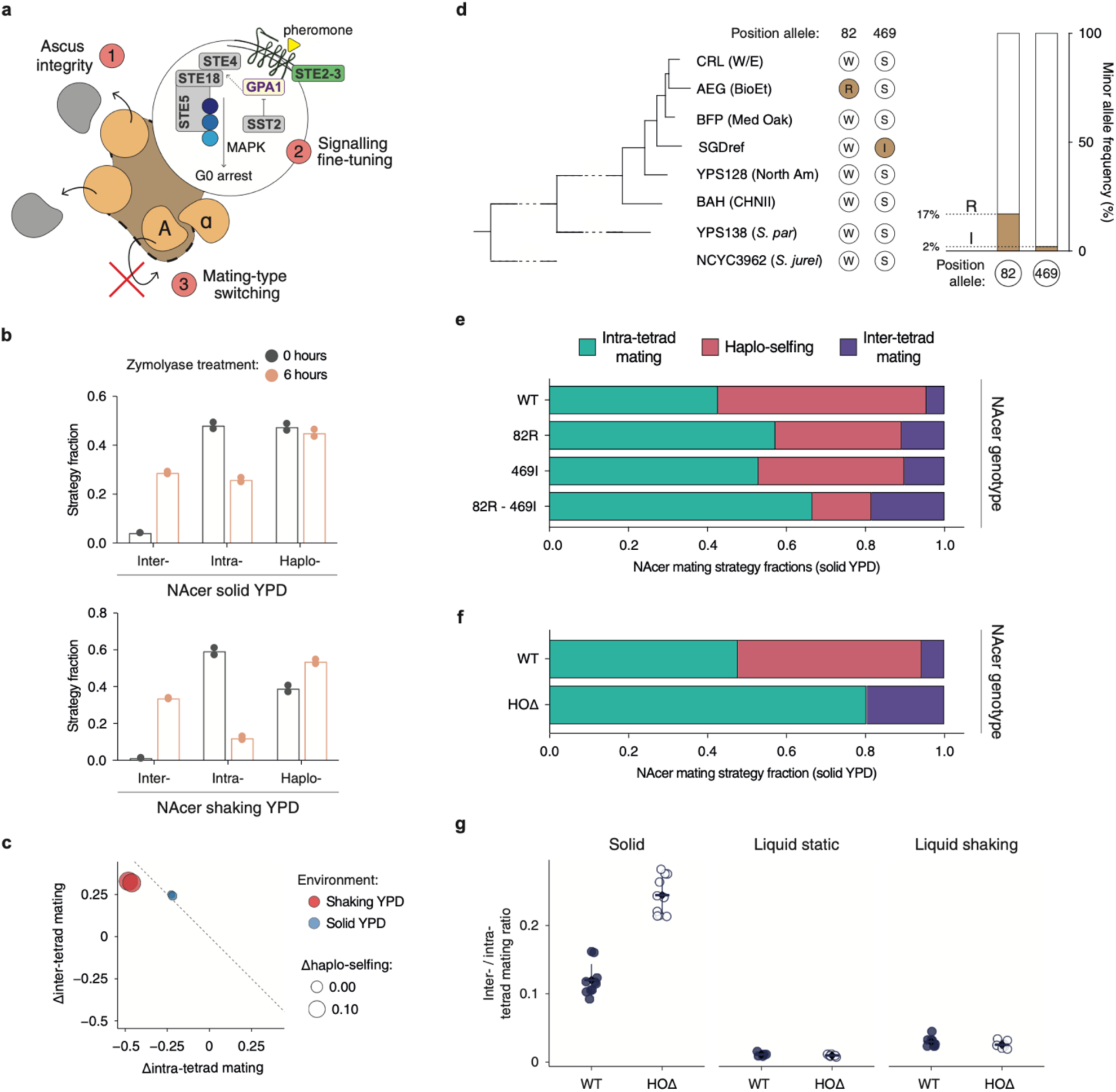
Genetic and ecological perturbations rewire mating outcomes. **a,** Cartoon illustrating three control layers acting at successive stages of the mating process: (1) physical release of spores which defines encounter geometry, (2) pheromone signaling which governs mating commitment, and (3) mating-type switching which determines access to haplo-selfing. **b**, Disruption of ascus integrity promotes inter-tetrad mating. Bar plots compare mating outcomes without (0 h) and after (6 h) zymolyase treatment in solid and shaking YPD environments (*N* = 2). Enzymatic digestion of the ascus wall shifted mating outcomes toward inter-tetrad fusion (Inter-), consistent with physical separation of sibling spores. Haplo-selfing (Haplo-) remained largely stable in structured conditions but increased modestly in shaking cultures, likely reflecting enhanced isolation of individual spores (Intra-corresponds to intra-tetrad mating). **c,** Changes in mating composition (Δintra- *vs* Δinter-; t₆-t₀) show a consistent loss of intra-tetrad and gain of inter-tetrad mating following ascus digestion. The diagonal denotes an equivalent exchange between the two modes, with deviations reflecting concomitant changes in haplo-selfing (circle size). **d,** Distribution of *GPA1* alleles across a collection of *S. cerevisiae*. Minor alleles (filled circles and filled bars) at both positions are rare within *S. cerevisiae* and not conserved across species (**Figure S5**), indicating that they represent the derived state. **e,** Mating outcomes in NAcer strains carrying individual (82R; 469I) or combined *GPA1* variants (82R-469I), compared to the respective wild-type version (W82; S469). Engineering these alleles shifts the balance of mating strategies, with the combined genotype producing a strong amplification of inter-tetrad mating under conditions that support pheromone signaling (solid YPD). This effect is largely absent in liquid static environment (**Figure S5A-B**), highlighting the interaction between signaling dynamics and environmental fluidity context. Bars represent mean mating fractions across independent replicates (*N = 7* in WT, *N = 6* in 82R, *N = 6* in 469I, and *N = 4* in 82R-469I). **f,** Removal of haplo-selfing through *HO* deletion redistributes mating outcomes towards both intra- and inter-tetrad mating. Bars represent mean mating fractions across replicates (*N = 10*). The increase in inter-tetrad mating upon *HO* deletion is maximal on solid (FC = 3.4X; **Table S5**). g, Relative balance between inter- and intra-tetrad mating in WT and *HOΔ* across the indicated environments. Points represent individual measurements and horizontal lines indicate mean ± SD. *HO* deletion significantly shifted the inter-/intra-ratio on solid medium (p-value = 2.31e-08), whereas differences in static and shaking cultures were not significant (Welch’s t-tests on log₂-transformed ratios).

In nature, ascus digestion in insect guts releases sibling spores and increases opportunities for outcrossing^23,32^. Thus, ascus rupture shifts mating opportunities away from siblings, with the physical environment determining the fate of released spores, mirroring its ecological role in facilitating outcrossing during insect dispersal^22^. To mimic this mechanism, we exposed sporulated asci to zymolyase, a β-1,3-glucanase enzyme that dissolves the ascus wall before germination. In both solid and shaking environments, inter-tetrad mating rose sharply over intra-tetrad mating, consistent with disruption of the spatial proximity between sibling spores (**Figure 2B**). By contrast, haplo-selfing remained stable on solid medium but increased modestly in shaking cultures, potentially reflecting higher isolation of individual spores after ascus degradation. To evaluate how mating modes trade off against one another, we compared net changes in intra- and inter-tetrad fractions following zymolyase treatment (Δinter vs Δintra; **Figure 2C**). On solid medium, the loss of intra-tetrad mating was almost entirely matched by increased inter-tetrad mating, whereas under shaking conditions it was partitioned between inter-tetrad mating and haplo-selfing.

Once spores are physically released, mating outcomes depend on their ability to sense and respond to potential partners. We therefore asked whether rewiring a single signaling node within the mating pathway could reshape the balance among mating strategies. *GPA1* encodes the Gα subunit that transduces pheromone-receptor activation to the downstream MAP kinase cascade, controlling cell-cycle arrest and mating-specific transcription (**Figure 2A**)^33,34^. We engineered two naturally occurring *GPA1* allelic variants previously shown to enhance mating efficiency (W82R and S469I) into our focal *Saccharomyces* NAcer background, which carries the major *GPA1* allele (W82/S469; **Figure 2D-E**; **Methods**). Both substitutions have been associated with increased pathway activation, by prolonging signal duration and up-regulating genes involved in mating response^35–37^. When expressed individually, each variant doubled inter-tetrad mating relative to the wild type (WT = 4.5% ± 0.5%; 82R = 10.9% ± 0.7%; 469I = 10.1% ± 1.3%), consistent with heightened pheromone responsiveness and an extended G1 arrest increasing partner encounter probability. Remarkably, combining both variants (82R-469I) increased inter-tetrad mating approximately 4-fold on solid medium (82R-469I = 18.5% ± 0.2%). This effect was highly environment-dependent and largely absent in liquid static cultures, where pheromone gradients are unstable (**Figure S5A-B; Table S2**). Contrary to what is observed in haploid strains^37^, the engineered lines displayed no detectable differences in growth rate (μmax) under standard rich-medium conditions, consistent with the repression of *GPA1* in diploid cells^33^ (**Figure S5C**). The combined genotype is extremely rare in natural populations, occurring in only eight strains among a collection of 3126 *S. cerevisiae,* with heterozygosity maintained at both variant positions (**Figure 2D**; **Figure S5D; Table S4**). This raises the possibility that the hyperresponsive *GPA1* genotypes may be selectively disfavored in fluctuating environments or in nutrient-limited niches, where prolonged signaling could delay proliferation or trigger maladaptive cell-cycle arrest.

More generally, enhanced signaling can increase mating propensity without guaranteeing cell fusion, as reproductive outcomes remain contingent on the availability and the timing of compatible partners. When partners are absent or delayed, cells can restore diploidy through haplo-selfing, introducing a final constraint on reproductive outcomes. At the broadest level, deletion of the endonuclease HO, the gene responsible for mating-type switching^38^, abolishes haplo-selfing entirely and redistributes mating outcomes between the two remaining routes^39^. In absolute numbers, removal of selfing predominantly favored intra-tetrad mating (**Figure 2F**). On solid medium, however, inter-tetrad mating increased proportionally more than intra-tetrad mating (FC = 3.4X *vs* FC = 1.7X, respectively; **Table S5)**. By contrast, in liquid cultures, *HOΔ* populations remained dominated by within-tetrad events (93.1% ± 2.8% in shaking, 94.4% ± 0.9% in static), consistent with physical clustering limiting access to alternative partners even when selfing is blocked. Accordingly, *HO* deletion significantly shifted the inter- to intra-tetrad mating ratio towards inter-tetrad mating on solid medium, whereas no significant shift was detected in either liquid environment (**Figure 2G**). These results suggest that removal of haplo-selfing can substantially increase the realized frequency of inter-tetrad mating when the physical environment permits the access to alternative partners. Consistent with this interpretation, recent population-scale analyses have associated loss of mating-type switching with increased heterozygosity and admixture in domesticated and mosaic heterothallic populations, suggesting that genetic constraints on selfing can increase opportunities for outcrossing^40,41^.

### Environmental context reshapes the mating landscape

The plasticity of mating outcomes across successive levels of reproductive control raises the question of how this flexibility responds to the environmental context in which mating occurs. Although the molecular and cellular mechanisms leading to yeast mating have been extensively dissected under laboratory conditions, how environmental cues are integrated by yeast cells to shape mating outcomes remains largely unexplored. When germination and mating occurred in media differing in nutrient complexity and affecting metabolism, the balance among mating modes was substantially reshaped (**Figure 3A**, **Figure S4** and **Table S2**). In Synthetic Oak Exudate (SOE)^42^, a moderately complex medium supplying readily metabolizable sugars and growth factors, the fraction of intra-tetrad mating strongly increased relative to YPD in the three backgrounds (NAcer = 64.0% ± 1% JSdist = 0.2; CHNIIcer = 46.9% ± 3% JSdist = 0.05; NApdx = 54.7% ± 7% JSdist = 0.36), with smaller rise in inter-tetrad mating proportions (NAcer = 8.7% ± 1.6%; CHNIIcer = 13.2% ± 2%; NApdx = 10.4% ± 1.4%). Despite persistent quantitative differences among genetic backgrounds, all exhibited the same directional shift towards the within-tetrad mating frequency inferred from population genomics^14^. This common response is compatible with a germination-synchrony model in which rapid and coordinated germination increases the probability of mating, by compressing mating opportunities into a narrower temporal window^43^. By contrast, YP-galactose medium shifted the mating landscape towards haplo-selfing (NAcer = 70.4% ± 4% JSdist = 0.24; CHNIIcer = 51.2% ± 3% JSdist = 0.07; NApdx = 91.3% ± 7% JSdist = 0.18) and increased inter-tetrad mating at the expense of intra-tetrad fusions in the two *S. cerevisiae* isolates (NAcer = 9.9% ± 3.3%; CHNIIcer = 14.5% ± 1.6%), consistent with delayed or asynchronous germination extending opportunity for mating-type switching and encounters with non-sibling gametes.

**Figure 3.**
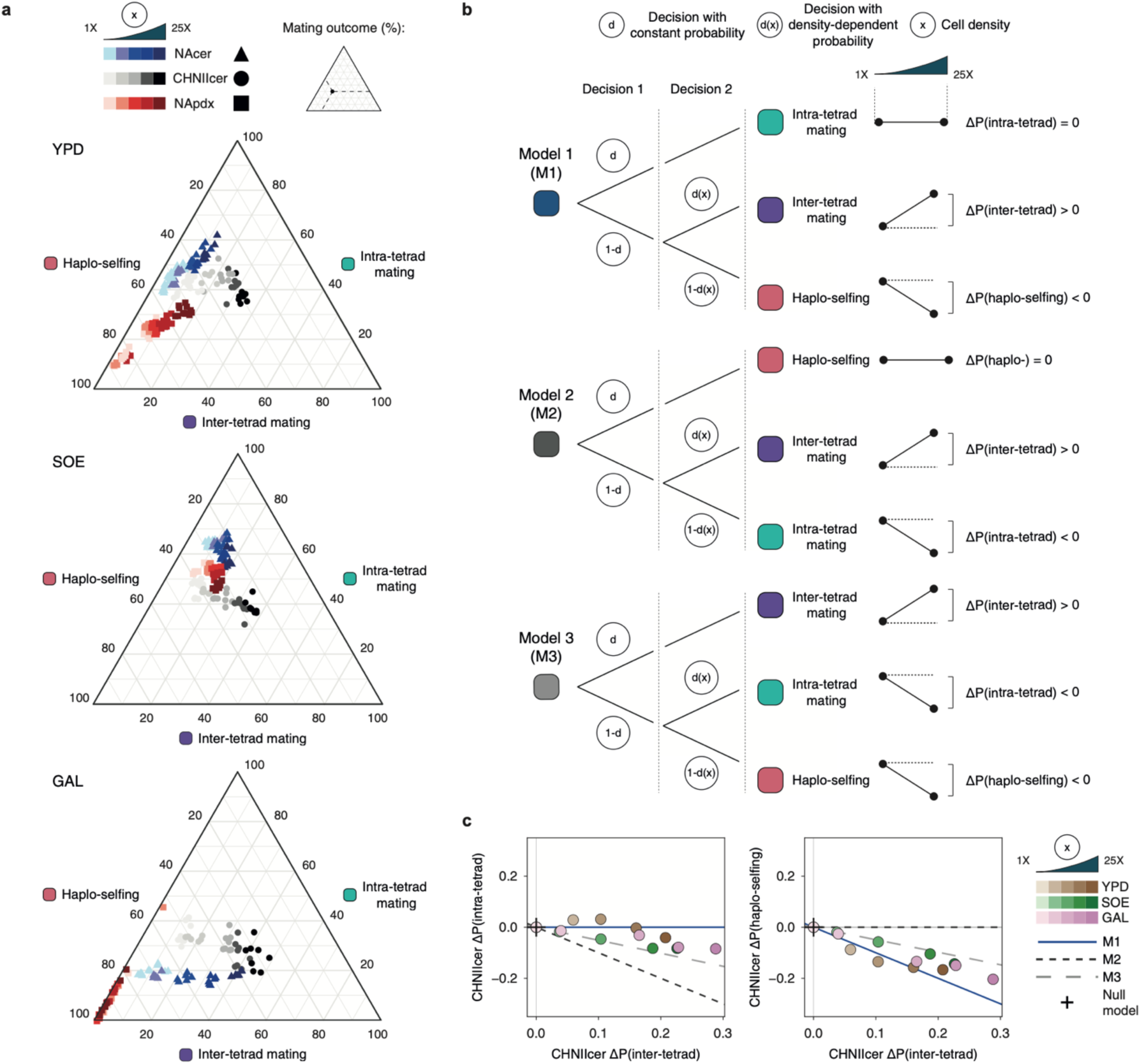
Environmental context and population density shape mating outcomes through hierarchical allocation of reproductive modes. **a,** Nutrient environment and population density jointly structure mating outcomes. Ternary plots show the distribution of intra-tetrad mating, haplo-selfing and inter-tetrad mating across media and cell densities in solid environment. Increasing density progressively shifts mating toward inter-tetrad events, whereas low-density conditions favor intra-tetrad mating and haplo-selfing. Each point represents an independent replicate. **b,** Conceptual models of hierarchical mating allocation. Schematic representation of alternative probabilistic hierarchies and their predictions across population densities. Model 1 (“*intra-first*”) predicts constant intra-tetrad mating; Model 2 (“*haplo-first*”) predicts constant haplo-selfing; Model 3 (“*inter-first*”) predicts a constant ratio between intra-tetrad mating and haplo-selfing. A non-hierarchical null model (“*no change*”) assumes equivalent mating routes. **c,** Density-dependent trade-offs reveal context-dependent mating hierarchies in CHNIIcer. Across the conditions shown, increasing density was consistently associated with declining haplo-selfing and increasing inter-tetrad mating, whereas intra-tetrad mating was less responsive, remaining approximately stable in YPD. The slopes of these trade-offs discriminate between alternative hierarchical scenarios: BIC-based model selection favored Model 1 in YPD, whereas Model 3 was better supported for solid SOE and GAL (**Appendix 2**).

To directly test how spatial encounter probability interacts with nutrient context, we next varied the population density at which spores germinate and mate, and quantified compositional mating outcomes in ternary space (**Figure 3A**). Because mating efficiency depends on the likelihood of contact between compatible gametes^11^, increasing cell density is expected to increase the opportunity for encounters between gametes from different tetrads. Consistent with this, higher density conditions promoted inter-tetrad mating, whereas sparse populations favored intra-tetrad mating and haplo-selfing. Notably, the magnitude of this shift was media and genotype dependent: combining nutritional dependency with the highest tetrad density fold change at inoculation (25 times higher than the standard one; **Methods**), both NAcer and CHNIIcer exhibited the largest density-driven increases in inter-tetrad mating in galactose (up to ∼8-fold and ∼4-fold, inter-NAcer = 40.8% ± 1%; inter-CHNIIcer = 43.3% ± 4.1%; **Table S2**). These shifts imply that nutrient-induced germination delays maximize the effect of spatial proximity. By contrast, inter-tetrad mating in NApdx showed a small opposite density dependency trend in galactose (inter-NApdx = 0.9% ± 0.2%), but responded markedly in YPD and SOE, suggesting genotype-specific sensitivity to carbon source or variation in germination traits (**Figure 3A; Figure S4; Table S2**).

To understand how mating modes respond to changes in population density across environments, we combined our quantitative measurements with predictions from a stochastic hierarchical mathematical model differing in the ordering of probabilistic mating decisions (**Figure 3B; Appendix 2**). In this framework, mating outcomes are represented as two sequential decision layers, where each event happens simultaneously. Spores allocate a fraction of mating interactions to one reproductive route, with the remaining fusion probability distributed among alternative outcomes. We compared three scenarios: (*i*) an “*intra-first*” model (Model 1), in which within tetrad mating is prioritized; (*ii*) a “*haplo-first*” model (Model 2), where auto-diploidization precedes any other interactions; (*iii*) an “*inter-first*” model (Model 3), assuming that between tetrad mating occupies the first layer. In all models, the probability of engaging in inter-tetrad mating increases with cell density, reflecting the higher encounter probability at higher cell crowding. We also included a non-hierarchical null model (“*no change*”) in which all probabilities are constant. Therefore, the models make distinct predictions as density changes: Model 1 predicts constant intra-tetrad mating, Model 2 constant haplo-selfing, and Model 3 a constant ratio between intra-tetrad mating and haplo-selfing.

Our data revealed a consistent decline in haplo-selfing and a rise in inter-tetrad mating as density increased across conditions (**Figure 3A**, **Table S2**), rejecting both Model 2 (“*haplo-first*”) and the null model. Overall, intra-tetrad mating was less responsive to density changes, remaining approximately stable in multiple conditions. Pairwise dependencies revealed a strong trade-off between haplo-selfing and inter-tetrad mating and a weaker trade-off between inter- and intra-tetrad mating (**Figure 3C**). Models 1 and Model 3 can both capture these relationships but predict distinct trade-off slopes, allowing them to be distinguished across conditions. Accordingly, BIC-based model selection showed that the preferred hierarchy varied across conditions (**Appendix 2).** Despite this variation, Model 1 provided the best overall description when conditions were considered jointly. Thus, reproductive outcomes are hierarchically organized, but their preferred hierarchy is not fixed across environments. Once sibling spores have fused, they are no longer available for subsequent mating events, such that density can influence only the distribution of the remaining reproductive outcomes.

### Within tetrad heterogeneity contributes to mixed mating

The hierarchical stochastic model derived from bulk cytometry measurements supported a hierarchical organization of reproductive outcomes. However, population-level measurements cannot determine directly whether spores within the same ascus realize uniform or divergent reproductive fates. To directly resolve this, we established a single-ascus complementary assay in which each individual tetrad was germinated separately, and the resulting colony analyzed by flow cytometry (**Figure 4A**). This design measures, for each ascus, the relative contribution of intra-tetrad diploids (green and red [G-R]), and haplo-selfed descendants (green [G-G] and red [R-R]), while excluding inter-tetrad events. Across all three backgrounds, individual tetrad-derived colonies frequently displayed substantial variation in the relative contributions of intra-tetrad mating, green haplo-selfed, and red haplo-selfed descendants (**Figure 4B, Table S7**). While population-level averages differed among strains, they often concealed extensive within-tetrad heterogeneity. Under identical environmental conditions, some asci produced colonies dominated by intra-tetrad mating-derived progeny, whereas others were enriched for haplo-selfed ones, with many generating mixed profiles. Importantly, the structure of this heterogeneity differed among backgrounds (**Figure S6**). In NAcer, [G-R] enriched tetrads were common, with the median [G-R] contribution (64.2%) exceeding the population average (50.2%), indicating that many tetrads were more strongly enriched for intra-tetrad mating than suggested by the mean composition alone. In contrast, NApdx exhibited highly skewed outcome distributions, with average [G-R] contribution (18.3%) driven by a minority of tetrads with exceptionally high [G-R] fractions, whereas the median tetrad contained only 0.2% of [G-R] descendants. However, and most importantly, NApdx is highly affected by lineage loss through gamete death. Consequently, the endpoint composition reflects both mating outcome and survival (**Table S8**). Instead, CHNIIcer displayed a distinct regime in which [G-G] and [R-R] descendants were broadly represented across tetrads despite substantial variation among individual asci. The strain population remains heterogeneous, but rather than being dominated by highly polarized tetrads, it appears to contain many tetrads in which multiple outcome classes coexist (**Figure S6**). Together, these observations indicate that spores within the same ascus diverge into distinct reproductive trajectories, and the population averages emerge from highly variable tetrad-level outcomes.

**Figure 4.**
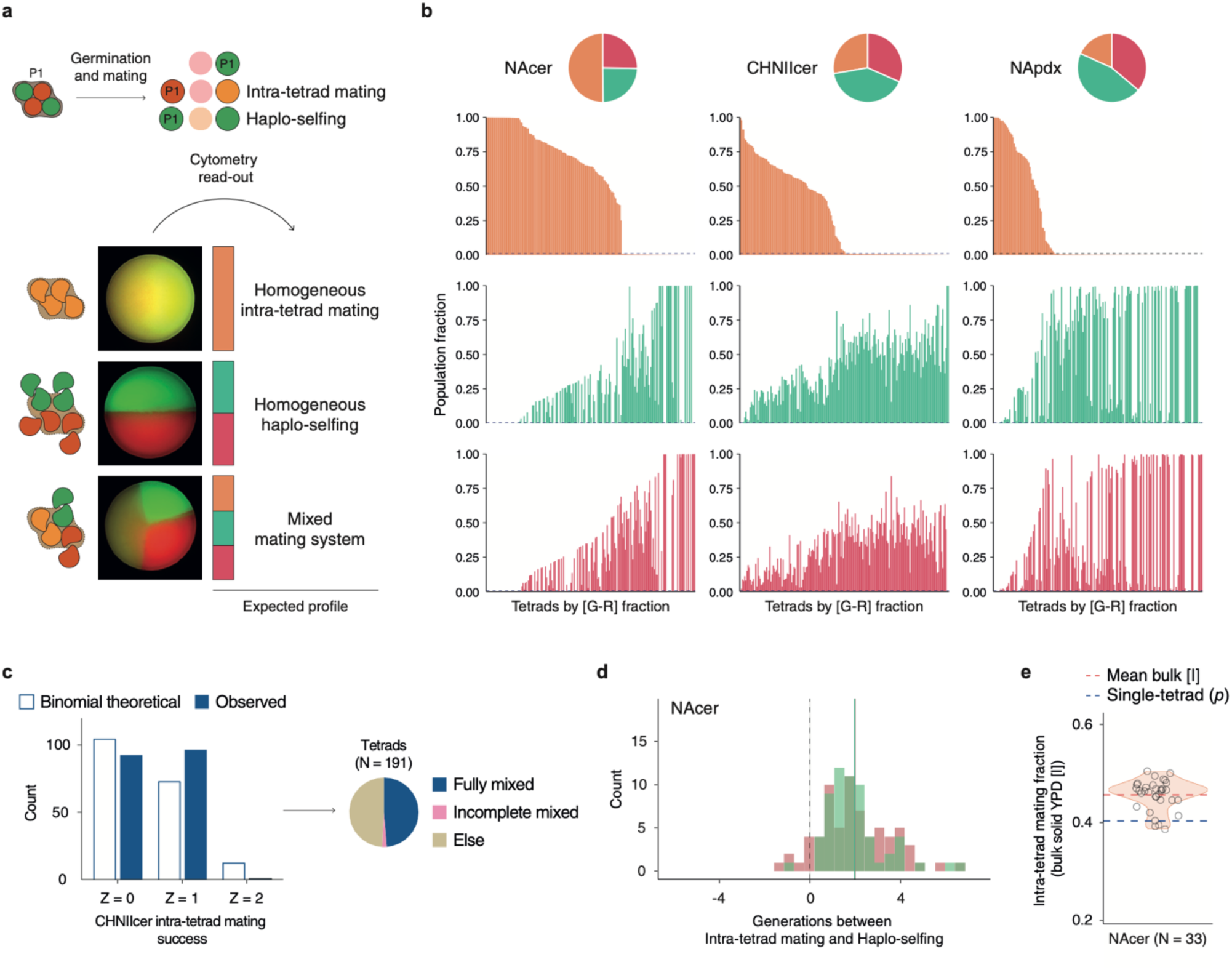
Tetrad-level heterogeneity in mating outcomes. **a,** Schematic of the single-tetrad assay. Individual tetrads were isolated, germinated on solid YPD, and allowed to form colonies. Flow cytometry was used to quantify the relative contributions of intra-tetrad mating [G-R], green [G-G] and red [R-R] haplo-selfed descendants. Representative stereomicroscopy images illustrate colonies exhibiting distinct patterns. Adjacent bar-plots depict expected profiles corresponding to colonies dominated by intra-tetrad mating, by haplo-selfing, or containing mixed populations. **b,** Composition of individual tetrad-derived colonies ranked by [G-R] contribution. Each bar represents a single tetrad and shows the fraction of descendants derived from intra-tetrad mating [G-R], green haplo-selfing [G-G], or red haplo-selfing [R-R] (NAcer, *N* = 196; CHNIIcer, *N* = 191; NApdx, *N* = 186). Dashed lines indicate detection thresholds for each class. Pie charts summarize the population-average composition across all analyzed tetrads, providing a population-level summary that contrasts with the heterogeneity observed among individual tetrads. **c**, Comparison between the observed distributions of intra-tetrad mating events per tetrad (Z), and the theoretical expectation under a simple binomial model in CHNIIcer (χ² = 19.87, p-value = 8.3e-06). The inset (pie-chart) further classifies individual tetrads according to their composition as fully mixed (all three classes detected), incomplete mixed (lacking one class), or else (only a single outcome detected), illustrating the relative contribution of each outcome to the observed distribution. **d**, Inferred temporal separation between intra-tetrad mating and haplo-selfing in NAcer. Analysis was restricted to Z = 1 tetrads (*N* = 96), which contain descendants from both reproductive modes. Histograms show the inferred number of generations by which green [G-G] and red [R-R] haplo-selfing lagged behind intra-tetrad mating. Generation 0 (vertical dashed line) corresponds to synchronous mating events, whereas colored vertical lines indicate the mean inferred delay (1.98 generations = cell divisions). **e**, Relationship between the probability of intra-tetrad mating estimated from the single tetrad assay (*p*) and its corresponding fraction in bulk solid 1X-YPD populations ([I]) in NAcer. The excess of [I] over *p* is consistent with the temporal separation inferred in panel **d**, as expected if these events occur earlier and therefore undergo more generations of expansion.

To dissect the structure of this heterogeneity, we inferred for each ascus the number of intra-tetrad mating events (denoted as “Z”, with Z ∈ {0,1,2}). These categories reflect the completed fusions between two compatible spore pairs: Z = 0 indicates fully haplo-selfing profile, Z = 2 indicates exclusive intra-tetrad mating, whereas Z = 1 represents a mixed mating system. We first estimated the probability that a compatible pair of gametes completes an intra-tetrad fusion (*p*), obtaining *p* = 26% (N = 191) for CHNIIcer, *p* = 40% (N = 196) for NAcer, and *p* = 16% (N = 186) for NApdx. We then compared the observed distribution of Z values with a binomial model parameterized by the probability of success *p* (**Methods**).

Only CHNIIcer significantly deviates from binomial expectation (χ² = 19.87, p-value < 0.001; **Figure 4C, Figure S6B**), likely driven by a marked excess of Z = 1 tetrads. Because accurate inference of Z requires the survival of all four spores, CHNIIcer, whose gametes viability is nearly complete (**Table S8**), is the only background in which this can be interpreted without ambiguity. Its deviation therefore potentially reflects real within-ascus diversification rather than artefacts of mortality or delayed germination.

In contrast, NAcer and NApdx did not significantly deviate from binomial predictions (χ² = 0.06 and χ² = 0.01 respectively, p-value = ns). Both strains exhibit higher gamete mortality, which can obscure coordinated behavior: loss or delay of one or more spores can distort the final color composition, thereby altering the inferred Z. For instance, the loss of all haplo-selfing derived lineages can cause a tetrad with a single intra-event (Z = 1) to be misclassified as Z = 2. Therefore, their binomial-like signatures likely represent a combination of independent outcomes and lineage mortality, rather than evidence against diversification.

Finally, for the subset of tetrads in which both lineages were detectable (Z = 1), we used their relative contribution to the final population to reconstruct the timing of lineage expansion from the first budding event (**Figure 4D**; **Methods**; **Appendix 3**). Across backgrounds, lineages ultimately contributing to intra-tetrad mating were consistently inferred to initiate growth several generations earlier than those producing haplo-selfed descendants (mean offsets: 1.18 in CHNIIcer, 1.98 in NAcer, and 1.82 in NApdx, **Figure S6**), revealing a systematic temporal asymmetry between the two reproductive fates. These differences in timing may explain why bulk assays measuring intra-tetrad fractions (mean [I]) exceed those predicted from single-tetrad probabilities (*p*) (**Figure 4E, Figure S6**). Together, within-tetrad heterogeneity is a fundamental feature of *Saccharomyces* reproductive behavior. This variability arises from a combination of intrinsic heterogeneity (as for CHNIIcer) and viability or timing (in NAcer and NApdx backgrounds). As a consequence, mixed mating at the population level may emerge not because different asci follow distinct strategies, but because individual spores within each ascus partition early reproductive fates.

### Temporal dynamics of germination and mating at single-cell resolution

To resolve how germination timing structures mating decisions, we developed a single-tetrad time-lapse microscopy pipeline tracking the complete reproductive path of each spore at single-cell resolution (**Methods**). For every tetrad, we annotated the time of ascus rupture and gamete release, the identity and timing of mating events (intra-tetrad or haplo-selfing), and any episodes of cell death (**Figure 5A**, **Video S1-2**). Time-lapse imaging revealed a strikingly ordered sequence of events: asci ruptured within a narrow temporal window, likely generating a transient phase in which spores were temporally aligned for subsequent mating interactions. In most tetrads, the first fusion was intra-tetrad, whereas the remaining spores remained on stand-by (and non-dividing) for several minutes despite physical proximity to potential compatible partners. This transient inactivity suggests that the initial intra-tetrad event may impose a local hierarchy, delaying alternative mating paths. Notably, early intra-tetrad matings did not constrain the remaining spores toward the same uniform outcome: later events were temporally dispersed, and frequently culminated in haplo-selfing, reinforcing the idea of asynchronous diversification rather than a fixed program (**Table S9**).

**Figure 5.**
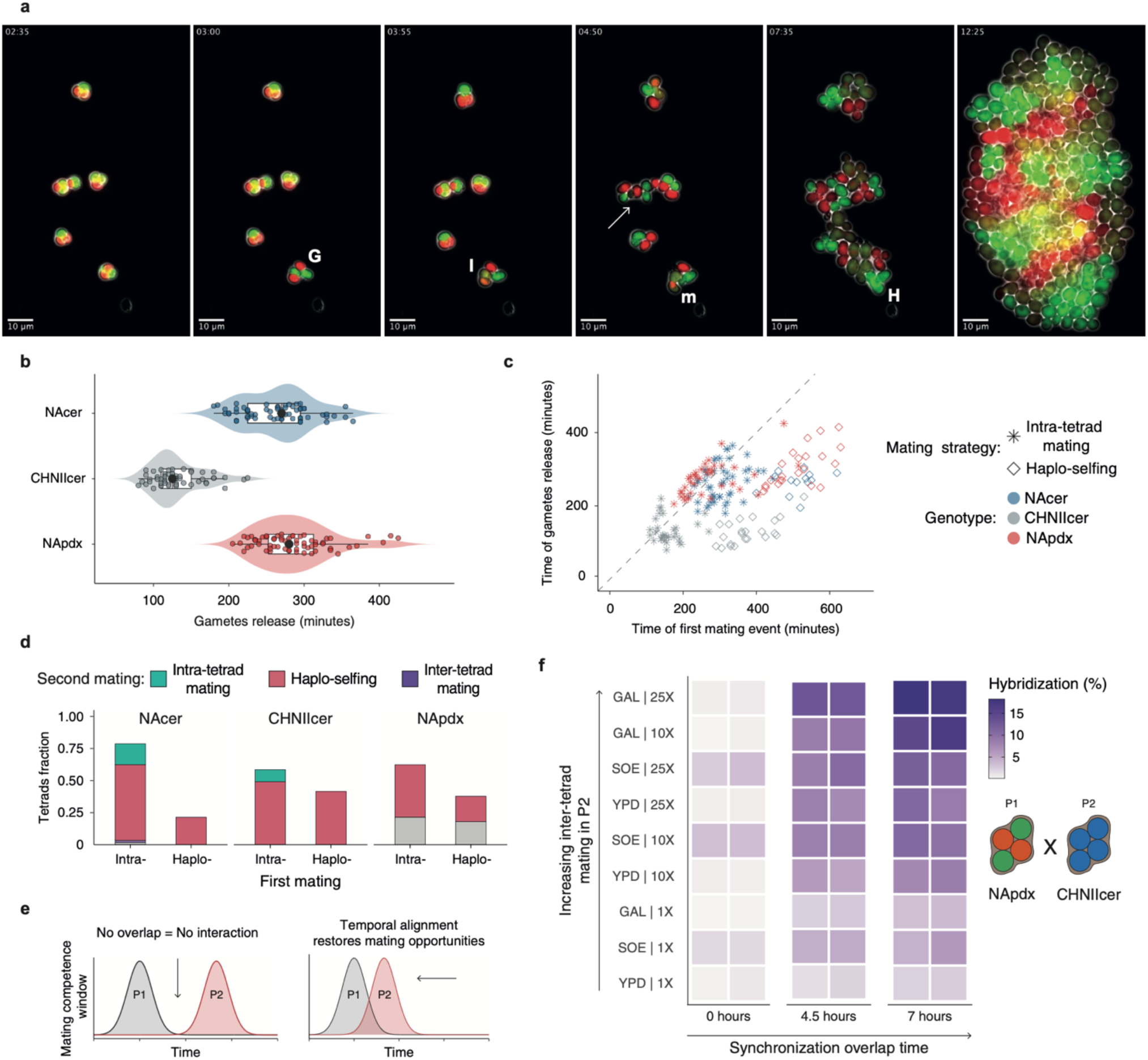
Germination dynamics define opportunities for mating and hybridization. **a,** Representative frames from single-spore time-lapse imaging (**Video S1-2**). Following ascus rupture and release of the first germinating spores (G), the earliest mating event typically occurs between siblings within the tetrad (I). Subsequent frames show continued spore release, coexistence of dormant and actively mating cells, cell death (arrow), mitotic proliferation of unmated haploid lineages (m), on-set of haplo-selfing (H), and the resulting mosaic colony composed of descendants originating from distinct mating trajectories. Scale bar: 10 μm. **b**, Timing of gamete release differs across genetic backgrounds. Distributions of gamete release times measured by single-spore time-lapse microscopy for NAcer (*N = 62*), CHNIIcer (*N = 53*), and NApdx (*N = 63*) tetrads. Violin plots show the full distribution, with individual asci shown as individual points. Central points within boxplots indicate the median. CHNIIcer exhibited earlier release, whereas NAcer and NApdx displayed delayed and broader temporal distributions. **c**, Temporal coupling between gamete release and first mating. Scatter plot showing the timing of the first mating event following ascus rupture for individual asci. Points are colored according to the genetic background (NAcer, CHNIIcer, and NApdx) and shaped according to mating strategy (intra-tetrad mating or haplo-selfing). The dashed line indicates zero delay between gamete release and mating. Intra-tetrad mating events remained more tightly coupled to germination than haplo-selfing events, which were delayed and exhibited greater temporal dispersion. **d**, The identity of the first mating event predicts subsequent mating outcomes. Stacked bars show the fraction of tetrads undergoing intra-tetrad mating or haplo-selfing as the first mating event across genetic backgrounds. Colors indicate the outcome of the second mating event within the same tetrad, including tetrads in which no second mating event was observed (in grey). Tetrads initiating with inter-tetrad mating or lacking a detectable first mating event were excluded from the analysis. Across backgrounds, the distribution of second mating outcomes depended on the identity of the first mating event, indicating significant dependence between sequential mating outcomes (Fisher’s exact test, p-value = 0.006; NAcer, *N* = 48 intra-tetrad and *N* = 13 haplo-selfing; CHNIIcer, *N* = 31 and *N* = 22; NApdx, *N* = 38 and *N* = 23). **e**, Conceptual model linking temporal overlap and hybridization potential. Under asynchronous germination, limited temporal overlap restricts opportunities for encounters between compatible gametes. Experimentally advancing germination of the slower population increases overlap between competence windows, restoring opportunities for inter-lineage mating. **f**, Temporal synchronization and ecological permissiveness jointly shape hybrid formation. Hybridization frequencies measured in CHNIIcer x NApdx mixes under asynchronous and experimentally synchronized germination regimes (4.5 and 7 hours in YPD; **Methods**) across ecological conditions differing in their capacity to promote inter-tetrad mating. Rows are ordered according to the inter-tetrad mating rate measured in isogenic CHNIIcer mixes under the corresponding conditions. Each tile represents an independent replicate, and color intensity indicates the fraction of hybrid diploids within the final diploid population.

To determine whether germination dynamics itself varies across genetic backgrounds, we quantified the timing of ascus rupture in each population (**Figure 5B**). CHNIIcer tetrads released their gametes earliest (median = 125 minutes) and showed the smallest temporal dispersion (SD = 33.8 minutes), producing a synchronized pulse of gamete availability. In contrast, NAcer and NApdx germinated later (medians = 270 and 280 minutes, respectively) and over broader temporal intervals (SD = 45.0 and 49.1 minutes), generating more temporally diffuse mating environments. These differences indicate that the temporal landscape of germination is itself genetically variable, extending previous observations of timing-dependent mate discrimination between *Saccharomyces* lineages^10^, likely modulating the balance between intra- and inter-tetrad mating in very dense populations. To determine how germination timing shapes mating routes, we next examined the temporal relationship between ascus rupture and the initiation of the first mating (**Figure 5C**). Across backgrounds, the timing of gamete release was significantly correlated with the timing of first mating (Spearman’s ρ = 0.45, p-value = 5.007e⁻10), indicating preserved temporal order between these processes. However, the dispersion around the 1:1 diagonal, together with a significant overall delay between germination and fusion (Wilcoxon signed-rank against zero delay, p-value < 2.2e-16), demonstrates that mating does not generally occur immediately upon gamete release but instead follows a post-germination decision window, during which reproductive fate remains unresolved. A two-way ANOVA revealed significant additive effects of mating route (p-value < 2e-16) and genotype (p-value = 0.000233) on the duration of this interval, with no evidence for a genotype by mating route interaction (p-value = 0.78). Thus, despite variation in absolute delay among genetic backgrounds, the germination-to-fusion interval strongly associated with the mating strategy (intra-tetrad mating *vs* haplo-selfing), with a consistent relationship across populations.

Consistent with this, intra-tetrad fusions occurred earlier and remained more tightly coupled to germination than haplo-selfing events, which were delayed and exhibited greater temporal dispersion. Tukey’s post hoc comparisons confirmed that haplo-selfing increased the germination to fusion delay by Δ = 209 min relative to intra-tetrad mating (95% CI: 191-227 min; p < 0.0001). Delays were shorter in NApdx compared to both NAcer (17–65 minutes smaller, 95% CI; p-value = 0.0003) and CHNIIcer (8–58 minutes smaller, 95% CI; p-value = 0.007), whereas NAcer and CHNIIcer did not differ significantly (p-value = 0.73). This temporal offset is consistent with the intrinsic generational constraint of haplo-selfing, which requires post-germination proliferation and mating-type switching before compatible descendants can fuse^44^. We next asked whether the identity of the first mating event biases subsequent decisions within the same tetrad (**Figure 5D**). Across backgrounds, intra-tetrad fusions dominated as the first mating event (NAcer = 78.7%, CHNIIcer = 58.5%, and NApdx = 62.3%), with haplo-selfing accounting for the remaining events. Conditional analysis of second mating outcomes revealed strong sequential bias: tetrads initiating with intra-tetrad fusion frequently proceeded to haplo-selfing (NAcer = 75%, CHNIIcer = 83.9%, and NApdx = 65.8%), whereas tetrads initiating with haplo-selfing consistently continued haplo-selfing (100%, 100%, 52.2% for NAcer, CHNIIcer, and NApdx respectively). Fisher’s exact test confirmed that first and second mating events are not independent (p-value = 0.006), demonstrating that reproductive outcomes are sequentially structured. This joint probability mapping illustrates a flexible hierarchy in which intra-tetrad mating predominates initially among the first events, while remaining spores can pursue alternative fates depending on both the initial outcome event and the genetic background. Notably, NApdx tetrads exhibited a higher frequency of incomplete second events, consistent with elevated spore mortality in this background and suggesting how viability intrinsically shapes the reproductive landscape (**Table S8**).

Finally, we asked whether genetically encoded differences in germination dynamics can generate a reversible form of temporal pre-zygotic isolation between divergent *Saccharomyces* lineages. Although reproductive isolation in yeasts is generally thought to arise predominantly through post-zygotic mechanisms^45^, asynchronous germination has previously been implicated in mate discrimination between *S. cerevisiae* and *S. paradoxus* spores^10^. To test whether restoring temporal overlap is sufficient to relax this constraint (**Figure 5E**), we quantified the raw hybridization rate as the fraction of yetSapphire-carrying diploids within the final diploid population derived from CHNIIcer[B-B] and NApdx[G-R] asci mixes (HYB fraction = f(mVenus– yetSapphire) + f(mCherry– yetSapphire); **Figure 5F**, **Table S10**). When tetrads from the fastest-germinating CHNIIcer background were mixed with those from the slowest NApdx background under standard conditions, hybrid diploids were nearly undetectable (<0.5%), even when population density and media composition were adjusted to maximize inter-tetrad mating in isogenic control cultures (**Figure 3A**). In these asynchronized mixes, CHNIIcer rapidly germinated and mated within the background, dominating endpoint cultures irrespective of the starting frequencies and effectively imposing a strong temporal barrier to hybrid formation. Strikingly, experimentally advancing *S. paradoxus* germination to align its mating-competence window with the Chinese background increased hybridization to ∼2–3%. Combining temporal synchronization with inter-tetrad permissive ecological conditions further elevated hybrid diploids to ∼18% of the diploid population, approaching the maximum yetSapphire-*color* inter-tetrad contribution observed in isogenic CHNIIcer mixtures (∼ 20.5% in GAL medium, 25X cell density; **Figure 3A**, **Table S2**). These results suggest that temporal mismatch constitutes a major constraint on hybrid formation but may be modulated by environmental conditions that alter germination dynamics and, consequently, mating opportunities between populations.

## Discussion

By combining quantitative population assays with single-cell measurements, we show that sexual reproduction in *Saccharomyces* operates as a plastic mixed mating system, in which reproductive fates emerge from probabilistic choices distributed across spatial and temporal scales rather than from a fixed program. In this framework, the relative contribution of the alternative mating routes varies with population features, genotype, and the environmental context. This extends beyond the traditional population genomic estimates for which intra-tetrad mating is inferred to dominate yeast reproduction. In *S. paradoxus*, the widely cited estimate of ∼94% intra-tetrad mating carries substantial uncertainty (95% C.I.: 41–99.9%) and is inherently sensitive to model assumptions about demographic and recombination processes^14^. Our direct measurements similarly identify intratetrad mating as a common reproductive route, although its contribution is neither exclusive nor uniformly dominant (**Figure 1E-F**). Instead, haplo-selfing and inter-tetrad mating can contribute substantially depending on genetic and environmental context. Population-genomic estimates capture the realized genetic consequences of mating in natural populations, whereas our measurements resolve the reproductive processes from which such patterns can emerge. Substantial capacity for inter-tetrad mating can therefore co-exist with rare genetic outcrossing, as its realization additionally requires genetically distinct compatible gametes to encounter in space and time. Our results support a view of yeast mating in which reproductive fate emerges from competition between alternative routes, whose accessibility changes dynamically following germination. In homothallic strains, intra-tetrad mating, haplo-selfing and outcrossing draw upon the same pool of mating-competent haploids^46^, which become sequentially accessible after gamete release. We further show that this competition is organized along complementary spatial and environmental axes, which modulate partner availabilities and the relative accessibility of alternative reproductive routes. Consistent with partner encounter being a key determinant of reproductive fate, spatial confinement on solid substrates enhanced inter-tetrad mating by maintaining physical proximity and by restricting pheromone gradients, whereas disruption of ascus integrity further shifted mating outcomes towards between-tetrad fusion (**Figure 2**). Consistent with this principle, insect-mediated dispersal can enhance outbreeding through ascus digestion, by releasing spores into mixed populations^22,23^.

Our results further identify temporal coordination as a complementary axis. Rather than directly favoring one mating route over another, environmental conditions additionally reshape the temporal window during which alternative routes remain accessible. Conditions promoting rapid and coordinated germination^43^ compress this temporal window, favoring within-tetrad mating, whereas delayed or asynchronous germination^47^ prolongs opportunities for mating-type switching and encounters with non-sibling gametes. Genetic regulation of pheromone signaling acts upon the same opportunity landscape. Manipulation of *GPA1* profoundly redistributed mating outcomes, increasing both inter- and intra-tetrad mating, consistent with quantitative variation in pathway activity modulating reproductive investment^36,37^. Our results extend this relationship from mating efficiency to realized mating mode, showing that modulation of pathway activity can alter the route through which diploidy is ultimately restored.

A simple stochastic model, with few parameters, further reveals that these population-level patterns are incompatible with equivalent, non-hierarchical mating routes and instead support preferred probabilistic hierarchies among reproductive modes whose organization shifts across conditions (**Figure 3**). Intra-tetrad mating remained comparatively less responsive to population density, whereas increasing density redistributed mating outcomes between haplo-selfing and inter-tetrad mating, consistent with their greater dependence on partner availability. Importantly, these hierarchies represent probabilistic preferences rather than obligatory sequences, allowing alternative routes to remain accessible within the same population. The model is intentionally agnostic to explicit time and therefore it cannot capture temporal differences in the realization of alternative mating routes. Such temporal differences may nevertheless influence endpoint measurements, with the relative enrichment of earlier-fusing lineages partly reflecting a proliferative head start rather than a greater probability of entering this route. Incorporating explicit timing into future models should refine quantitative predictions while preserving the central inference that reproductive outcomes are hierarchically organized.

Independent single-tetrad analyses support the predicted ordering of within-tetrad events and unfold their dynamic realization at the level of individual gametes (**Figure 4**). These analyses also reveal a key feature inaccessible to population-level measurements: mixed mating can arise through reproductive diversification within the same tetrad. Such heterogeneity may partly reflect how individual spores respond to their local context, as pre-established spore polarity can be re-oriented by mating cues^48^. Regardless of its underlying mechanism, this diversification challenges the conventional interpretation of haplo-selfing. Haplo-selfing has often been interpreted as a fallback route engaged only when direct sibling mating is prevented, for example by spore unviability or physical separation^9,20,39^. Instead, our data identify haplo-selfing as an intrinsic competitor within the reproductive landscape, rather than a passive consequence of failed encounters. McClure et al. (2018)^49^ previously showed that sibling spores can differ in their post-germination behavior and proposed delayed mating within mixed-mating-type microcolonies. By following individual gametes, we show that this heterogeneity translates into distinct realized reproductive routes within individual asci. Moreover, this within-ascus diversification is structured by a systematic temporal asymmetry between reproductive fates. Across genetic backgrounds, intra-tetrad diploids consistently initiated growth earlier than haplo-selfed descendants. Early intra-tetrad fusions therefore acquire a proliferative head start while other meiotic products are still progressing through the mitotic and mating-type-switching steps required for haplo-selfing, potentially biasing endpoint measurements towards faster reproductive routes. Importantly, the identity of the first mating event influenced subsequent outcomes within the same tetrad shaping but not uniquely determining, the fate of the remaining sibling gametes (**Figure 5**). Different products of the same meiosis can consequently realize distinct reproductive paths towards intra-tetrad mating, haplo-selfing or, when appropriate partners are encountered, outcrossing. Consequently, the ascus should not be regarded as a uniformly behaving reproductive unit. Despite sharing ancestry and an immediate environment, its four gametes can frequently diverge in reproductive fate.

Our time-resolved imaging at the single-tetrad level provides a mechanistic view of how mating unfold in real time, revealing that physical proximity between compatible gametes alone does not guarantee immediate fusion. Mating in *Saccharomyces* follows a structured, though stochastic, temporal sequence and is embedded within a broader post-germination window that varies across genetic backgrounds, generating genotype specific mating landscapes. While our imaging experiments were conducted in simplified controlled microcosms, natural environments are expected to amplify these dynamics through spatial heterogeneity and fluctuating conditions. Future work should therefore quantify how mating-competence windows vary across environmental gradients and assess whether selection acts on their timing and duration. Finally, while our manual annotation of time-lapse data yielded robust and reproducible insights here, scaling these analyses will require automated image-analysis pipelines^50,51^.

The causal importance of temporal overlap is particularly evident in our hybridization experiments (**Figure 5**). Under standard conditions, hybridization between the fastest (CHNIIcer) and the slowest (NApdx) germinating strains was nearly undetectable, consistent with previous evidence that differences in germination timing can contribute to prezygotic reproductive isolation^10^. Experimentally aligning mating competence windows was sufficient to reverse temporal reproductive isolation, with a stronger effect under conditions that maximize inter-tetrad encounters (**Figure 3A**). Crucially, the critical variable governing mating outcomes is not strict synchrony per se, but the extent and the duration of temporal overlap. Germination asynchrony can either suppress or facilitate hybridization, depending on whether it fragments or extends mating opportunities, respectively: when germination is excessively staggered, between-tetrad mating opportunities are fragmented and hybridization is suppressed; in contrast, population-level asynchrony can extend the period over which compatible gametes coexist (**Figure 6**). Germination timing therefore does not simply accelerate or delay mating but quantitatively structures the network of mating opportunities. The reversibility of temporal reproductive isolation further shows that reproductive outcome can be modulated without altering genetic compatibility. Manipulating temporal coordination may thus provide a new lever for applications in yeast breeding and strain engineering^52^.

**Figure 6.**
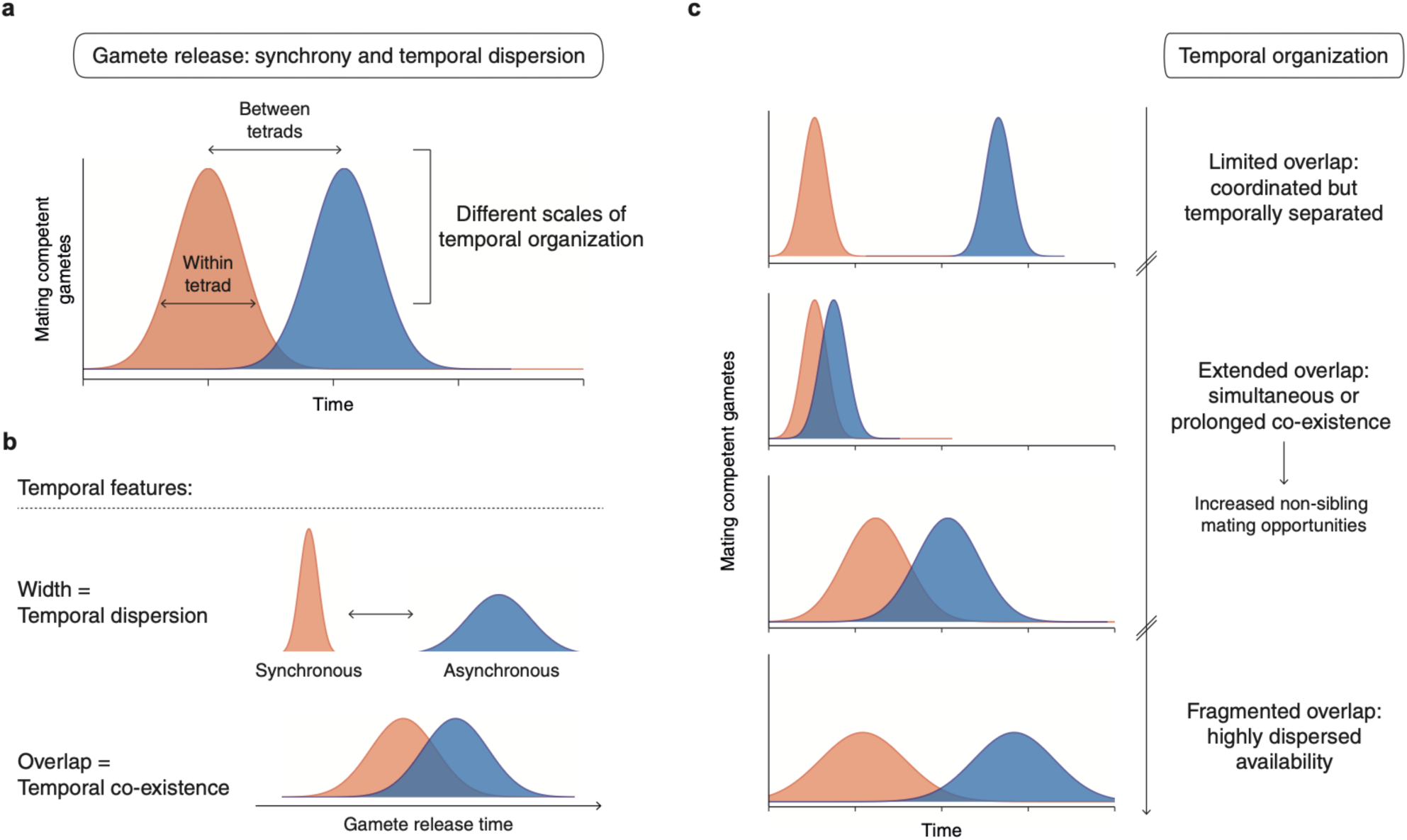
Temporal organization shapes mating opportunities. **a,** Gamete availability is shaped by temporal variation operating at different scales (heterogeneity among siblings and/or between tetrads). b, Temporal features underlying this organization. Curve width captures the degree of temporal dispersion in gamete release, whereas the overlap between distributions reflects temporal coexistence of mating-competence windows. c, Combinations of synchrony and dispersion generate distinct windows of mating opportunity. Separated pulses limit non-sibling mating, whereas overlapping distributions create concentrated or prolonged periods of gametes coexistence. At greater temporal dispersion, reduced overlap can fragment these opportunities.

These cellular dynamics have direct evolutionary implications, as selfing, intra-tetrad and outcrossing differ fundamentally in their consequences for genetic variation by controlling levels of inbreeding and outbreeding^39^. Even relatively rare outcrossing events can introduce genetic variation that is inaccessible through selfing. Population genomics analyses reveal extensive population structure and admixture during *Saccharomyces* diversification^15,26^. Moreover, multiple *S. paradoxus* introgressed genomic regions have been described across *S. cerevisiae* populations, indicating recurrent secondary contact^17,18^. Similarly, hybridization between divergent genomes can generate novel combinations of parental traits and facilitate adaptation to new ecological niches in domesticated yeasts^53,54^. Mechanisms that preserve even transient opportunities for mating with genetically distinct partners may therefore have evolutionary consequences disproportionate to the frequency of outcrossing itself, by providing access to genetic combinations unavailable through selfing.

Finally, diversification of reproductive fates within individual asci raises the possibility of reproductive bet-hedging. Microbial bet-hedging has largely been associated with phenotypic diversification within clonal populations, whereby genetically similar cells adopt alternative states that distribute risk across uncertain environments^55–58^. Here, by contrast, diversification occurs among sibling products of a single meiosis and affects their route to reproductive success. Distributing gametes across alternative reproductive trajectories could reduce the risk that an entire meiotic cohort experiences reproductive failure when mating opportunities are uncertain. Importantly, progeny generated through different routes showed comparable fitness under our experimental conditions (**Figure S1E**), indicating that diversification of reproductive access does not require detectable fitness differences. This resembles individual bet-hedging, in which diversification among descendants distributes risk across alternative phenotypic outcomes^59^ but extends this principle to reproductive fate. A related logic operates in multicellular organisms, where offspring can diversify their response to future conditions, for example through variation in seed dormancy and germination in plants or egg diapause and hatching in animals^60,61^. Testing this hypothesis will require determining whether diversified tetrads achieve higher long-term reproductive success than cohorts constrained to a single mating route across environments that differentially favor sibling mating, haplo-selfing or outcrossing.

More broadly, our findings support a view of yeast reproduction in which meiosis and mating are not obligatorily linked. Meiosis establishes a cohort of potential gametes whose eventual reproductive fate remains responsive to the conditions encountered after germination. Whether this organization can be fully explained by the interplay between pre-existing spore polarity and spatial state^48^, germination and cell-cycle progression, pheromone signaling, and partner availability, or involves additional regulatory layers governing reproductive strategy, remains an important mechanistic question. In this context, the ascus may function as a spatial and temporal buffer, favoring sibling mating without fully determining reproductive fate, while allowing alternative routes to emerge as environmental conditions and partner availability change. Such flexibility may reconcile reproductive assurance through selfing with opportunities for genetic diversification through outbreeding. Rather than being determined at meiosis, reproductive strategy remains responsive during a critical post-germination window, in which genotype, environment, space and time jointly shape who ultimately mates with whom.

## Supplementary Figures

**Figure S1.**
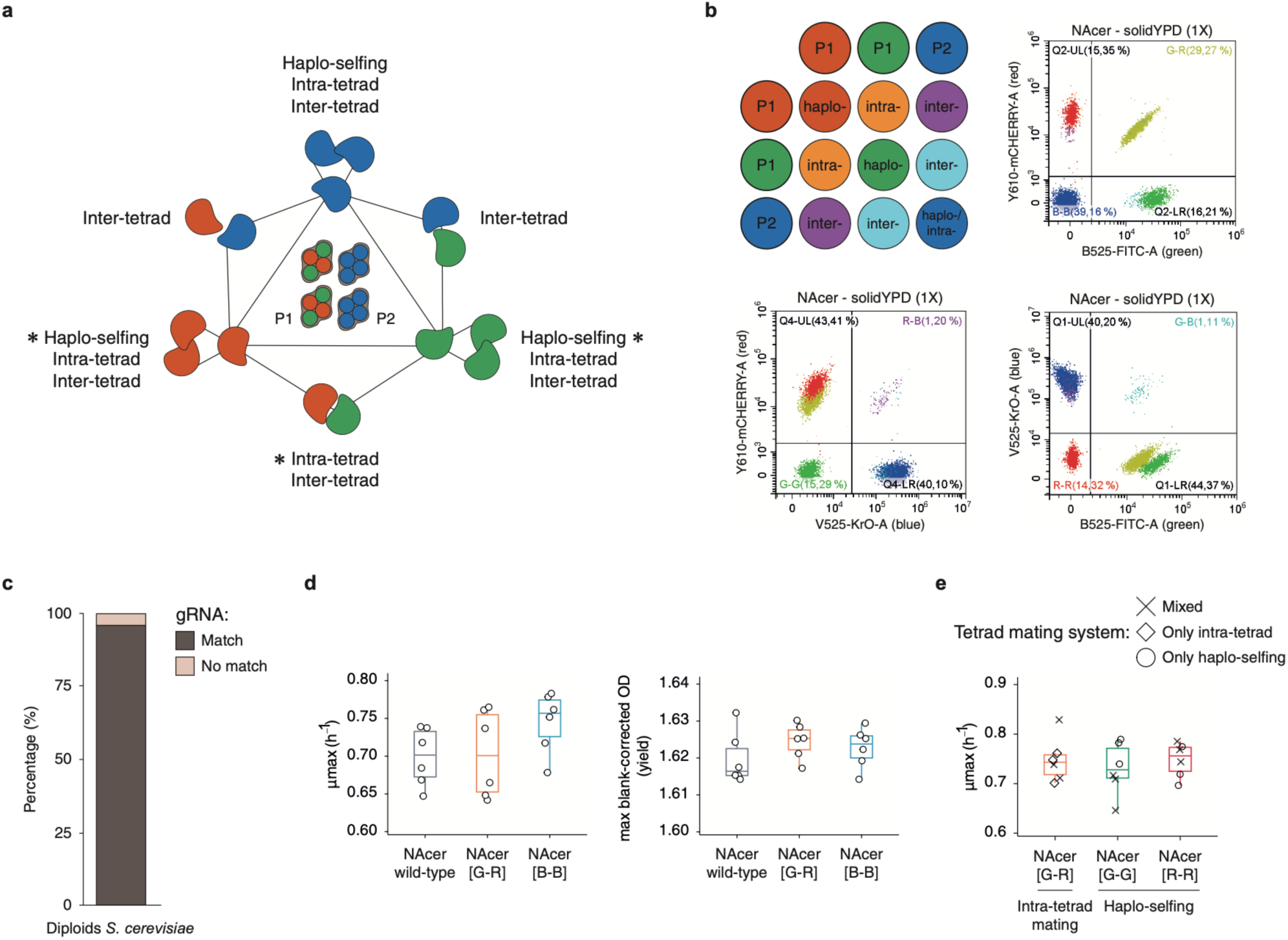
Experimental architecture and analytical workflow of gaMATe. **a**, Conceptual design of the system. Each color combination defines a distinct fluorescence signature corresponding to a specific mating outcome. Asterisks indicate the primary mating outcome associated with each color combination. Alternative biological scenarios can produce the same profile, for example as a result of recombination, and require an additional correction step as described in **Appendix 1**. **b**, Analytical workflow of the multicolor flow cytometry assay for interpreting pairwise (P1xP2 = asci mixes) mating assays. Representative plots illustrating the gating strategy used to assign events to each color combination corresponding to a specific mating outcome. Gates defining each population are indicated in the cartoon, together with the corresponding mating outcome inferred from each fluorescence profile. **c**, Predicted applicability of gaMATe across *S. cerevisiae*. The proportion of strains with complete gRNA sequence matching is shown, indicating that the system is compatible with ∼96% of the known *S. cerevisiae* genetic diversity^15^ (**Methods**). **d-e**, Growth phenotypes across genotypes show neutrality of the system. Boxplots represent maximum specific growth rate (μmax) and maximum blank-corrected OD (yield) for the indicated genotypes. Points represent individual measurements from independent replicates (*N* = 6), and center lines indicate medians. In **e**, symbols indicate strains derived from tetrads with different mating system profiles (mixed, only intra-, or only haplo-, Figure 4A). No significant differences between genotypes were detected by one-way ANOVA or Kruskal-Wallis tests.

**Figure S2.**
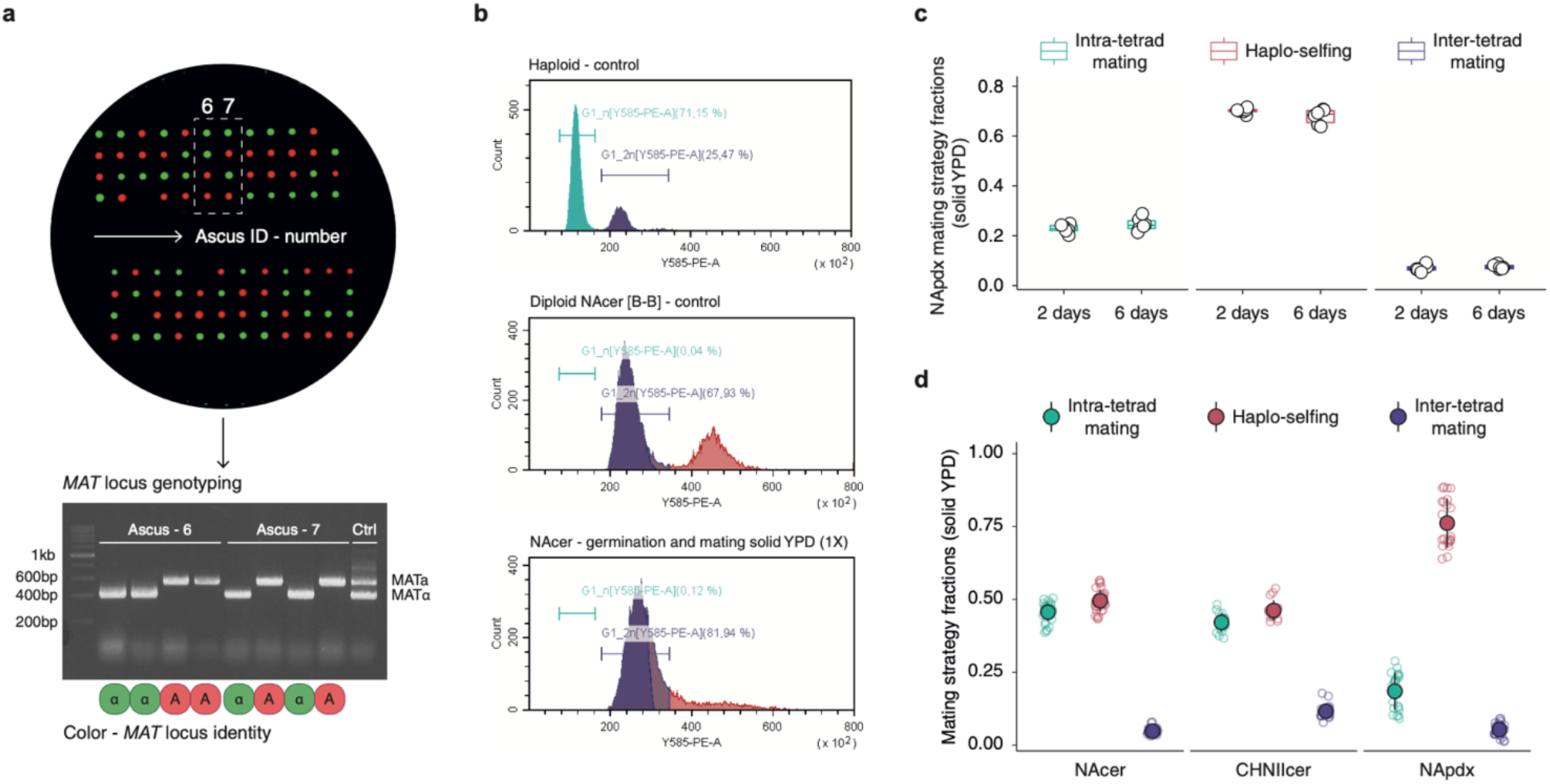
Experimental validation and robustness of gaMATe. **a**, PCR validation of *MAT* locus and fluorescence genotype linkage following meiosis. An *HO* deleted NAcer strain was sporulated and individual asci dissected by micromanipulation. Colony PCR on resulting colonies was used to verify the association between mating type and fluorescent reporter inheritance. The experimentally observed recombination rate (∼0.5%) was consistent with theoretical expectations (**Methods**). **b**, Verification of ploidy following germination and mating using propidium iodide (PI) staining as described in^15^. Representative DNA-content histograms obtained by flow cytometry for haploid and diploid controls, together with a representative mating culture, demonstrate the predominance of diploid cells following germination and mating. Quantification of residual G1_n events is provided in **Table S6**. **c**, Robustness of inferred mating strategy distributions to endpoint sampling. In NApdx, mating outcomes quantified after 2 or 6 days of incubation yielded comparable estimates, indicating that the inferred compositional distributions are stable across the tested sampling interval. Points represent individual measurements (*N* = 6), and center lines indicate medians. **d**, Replicate-level distributions of mating strategy fractions across genotypes. Individual points represent replicates across multiple independent experiments (NAcer, *N* = 33; CHNIIcer, *N* = 16; NApdx, *N* = 22), while large, filled circles denote means and error bars indicate SD. The distributions demonstrate the reproducibility of genotype-associated mating profiles across experiments while capturing the extent of within genotype variability.

**Figure S3.**
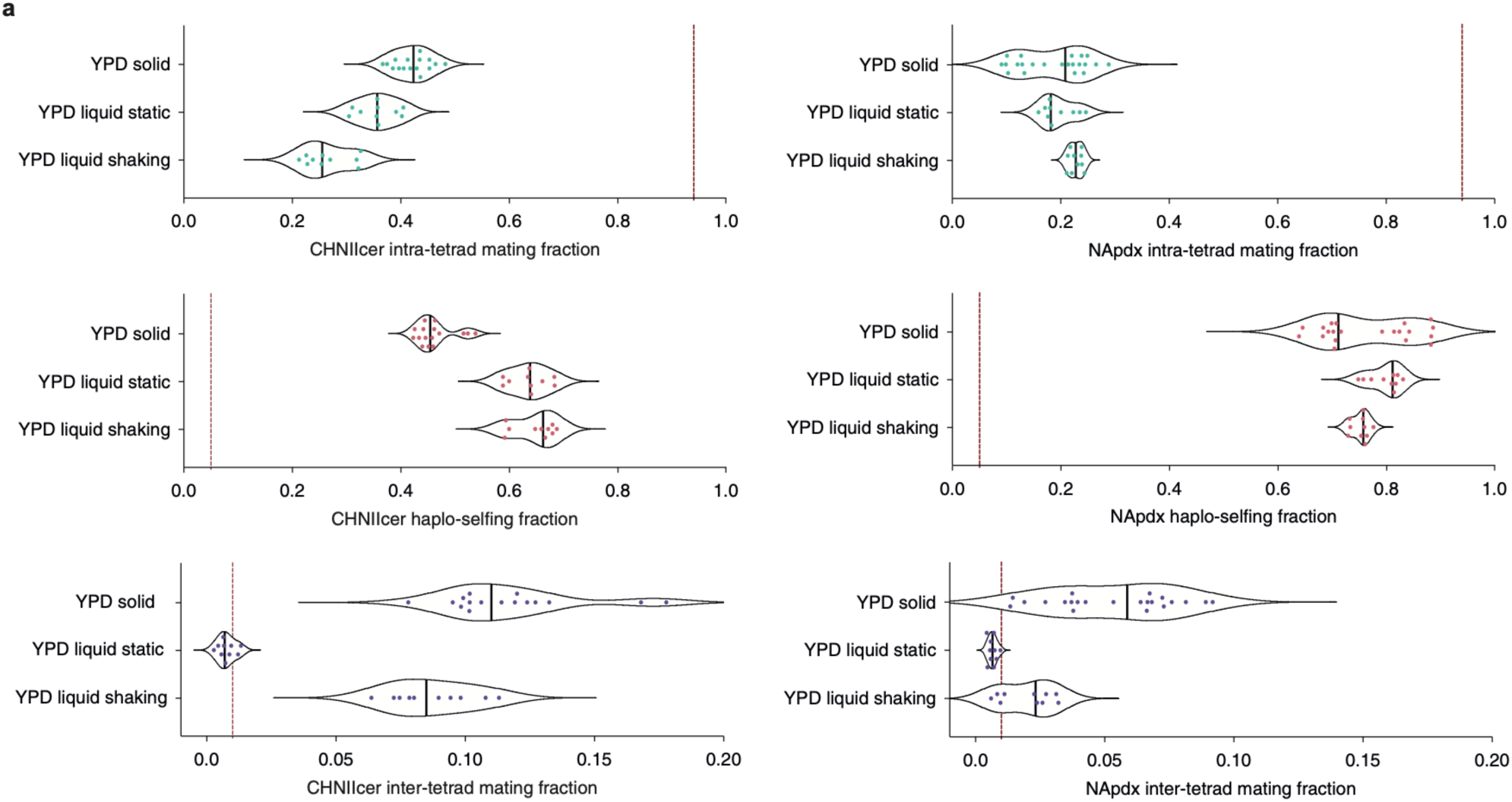
Effect of the physical environment on mating outcomes in additional genotypes. **a**, Violin plots showing the distributions of intra-tetrad mating, haplo-selfing and inter-tetrad mating fractions measured in CHNIIcer (on the left) and NApdx (on the right) under different physical conditions (solid agar, static liquid and shaking liquid YPD). Each point represents an independent replicate (*N* = 16 (CHNIIcer), *N* = 22 (NApdx) in solid YPD; *N* = 10 (CHNIIcer and NApdx) in both static and shaking liquid YPD). Central lines indicate medians. The vertical dashed line denotes the mating frequency inferred from population genomic analyses^14^.

**Figure S4.**
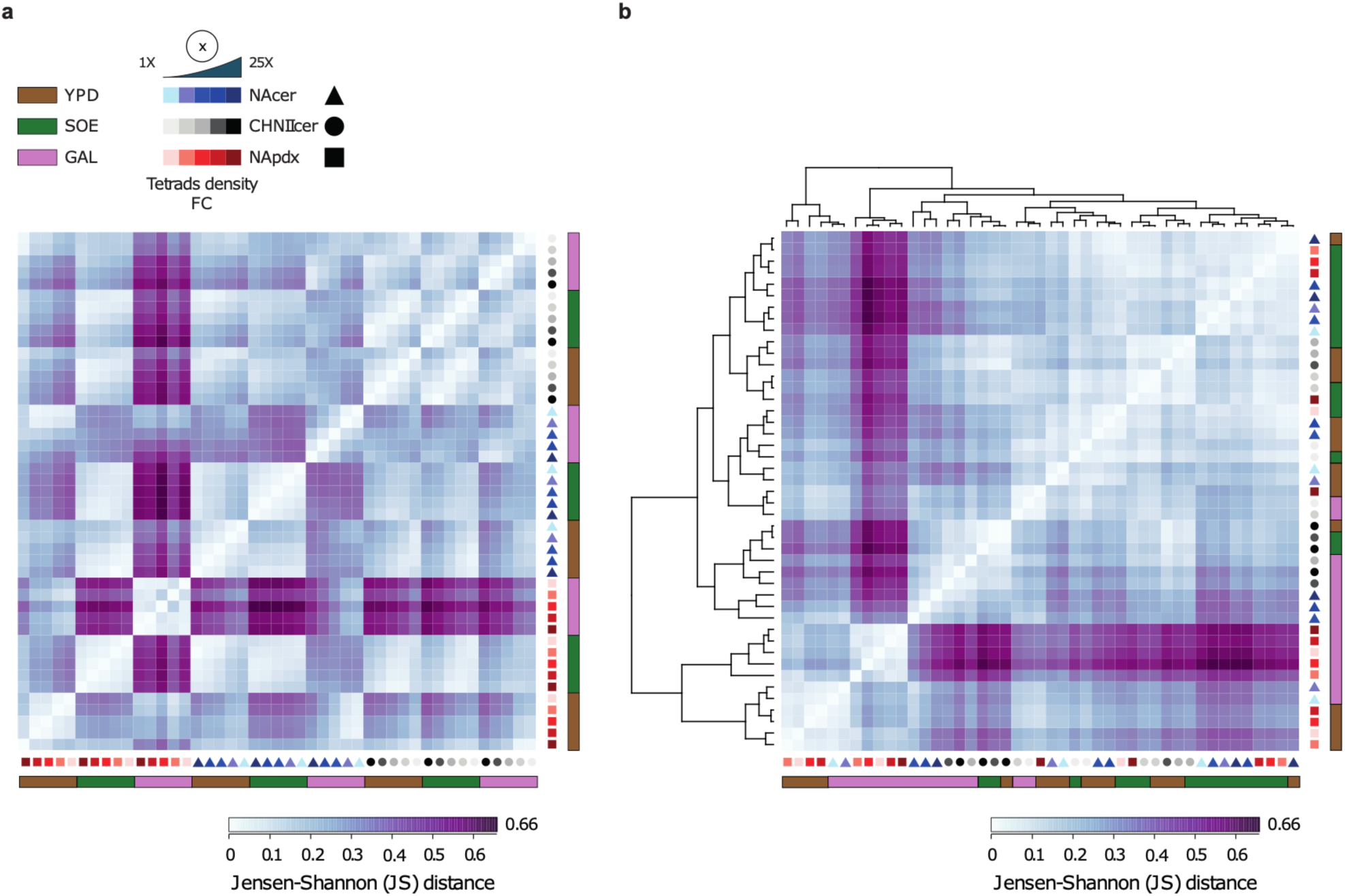
Similarity of mating strategy distributions across experimental conditions. **a**, Heatmap showing pairwise Jensen-Shannon distances (JSdist) between mating patterns across the different experimental conditions tested in solid environment. Distances were calculated as the square root of the Jensen-Shannon divergence, with smaller values indicating greater similarity. Experimental conditions were ordered according to strain, germination medium and cell concentration to facilitate visual comparison of the contribution of each factor. Symbols denote strain and cell concentrations, and colored bars indicate the medium. **b**, The same distance matrix shown in **a**, with experimental conditions ordered by hierarchical clustering. Hierarchical clustering revealed grouping of GAL and SOE media, whereas YPD conditions are distributed across multiple branches. Samples from NApdx in GAL formed two distinct clusters, highlighting higher separation from other experimental conditions.

**Figure S5.**
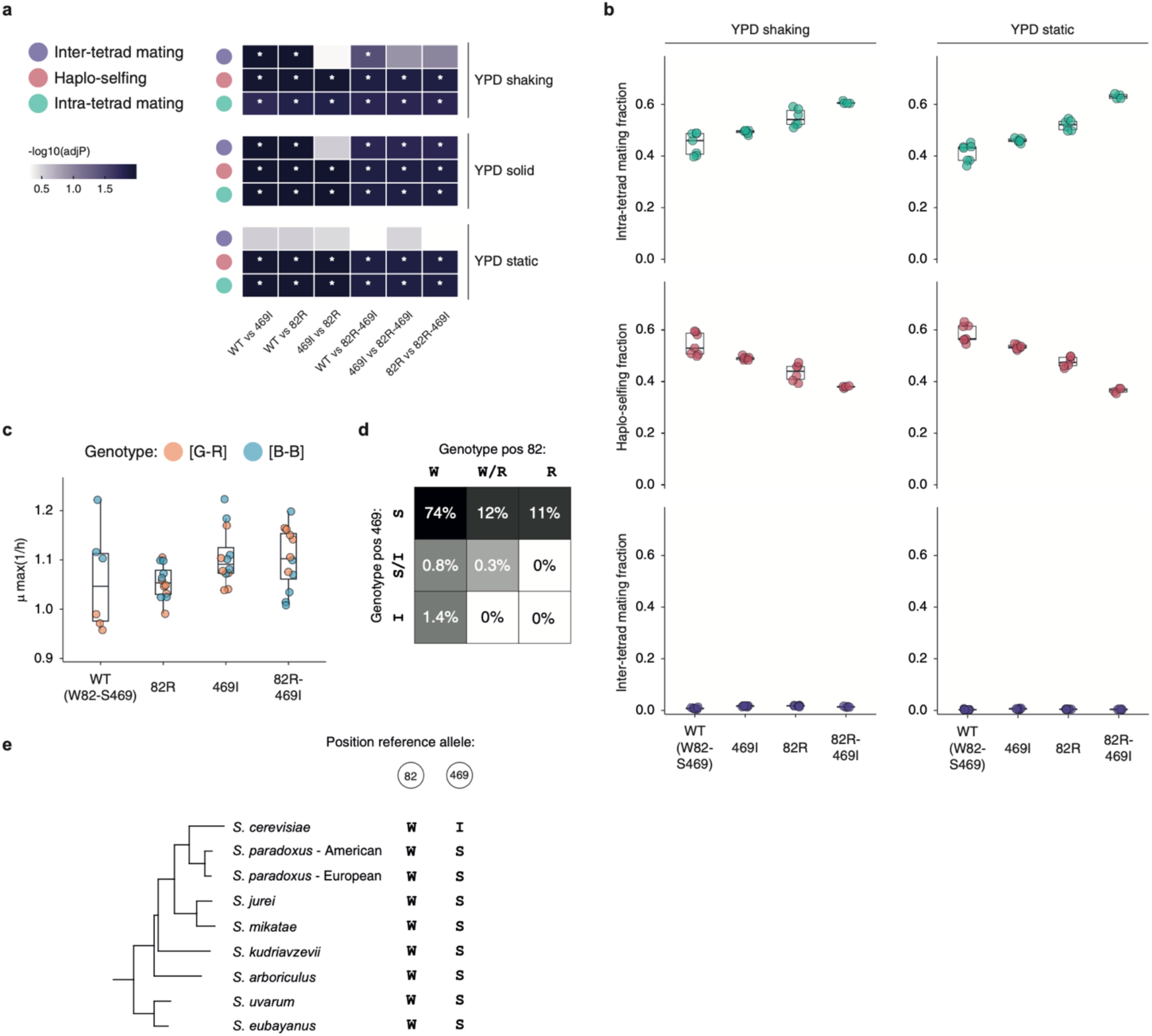
Context-dependent effects of GPA1 variants on mating outcomes. **a**, Statistical comparison of genotype dependent differences in mating strategy composition across environmental settings. The heatmap shows Benjamini–Hochberg (BH) adjusted p-values from pairwise Wilcoxon rank-sum tests comparing *GPA1* genotypes independently for each mating strategy and experimental condition. Color intensity represents −log10(adjP), with darker cells indicating stronger evidence for genotype-dependent differences, and asterisks denote statistically significant comparisons after correction. **b**, Box-plots show the distributions of intra-tetrad mating, haplo-selfing and inter-tetrad mating fractions measured across the indicated *GPA1* genotypes under static liquid and shaken liquid YPD conditions. Individual points represent independent replicates (*N = 7* in WT, *N = 6* in 82R, *N = 6* in 469I, and *N = 4* in 82R-469I) whereas colors denote mating strategy. Statistical comparisons are summarized in **a**. **c**, Growth phenotype of *GPA1*-engineered diploid strains. Boxplots show the maximum specific growth rate (μmax) across the indicated genotypes. Points represent individual replicates (*N = 6* in WT, *N = 12* in 82R, *N = 12* in 469I, and *N = 12* in 82R-469I) and are colored according to the fluorescent reporter genotype. No significant differences in μmax were detected under the conditions tested. **d**, Distribution of *GPA1* alleles at positions 82 and 469 across a collection of 3126 *S. cerevisiae* isolates. Heatmap colors and percentages indicate the proportion of strains harboring each allelic combination. The combined mutant genotype is rare among the surveyed natural isolates, possibly occurring only as double heterozygotes (0.3%). e, *GPA1* alleles within *Saccharomyces* genus. Minor alleles at both positions are not conserved across species, consistent with them representing the derived state.

**Figure S6.**
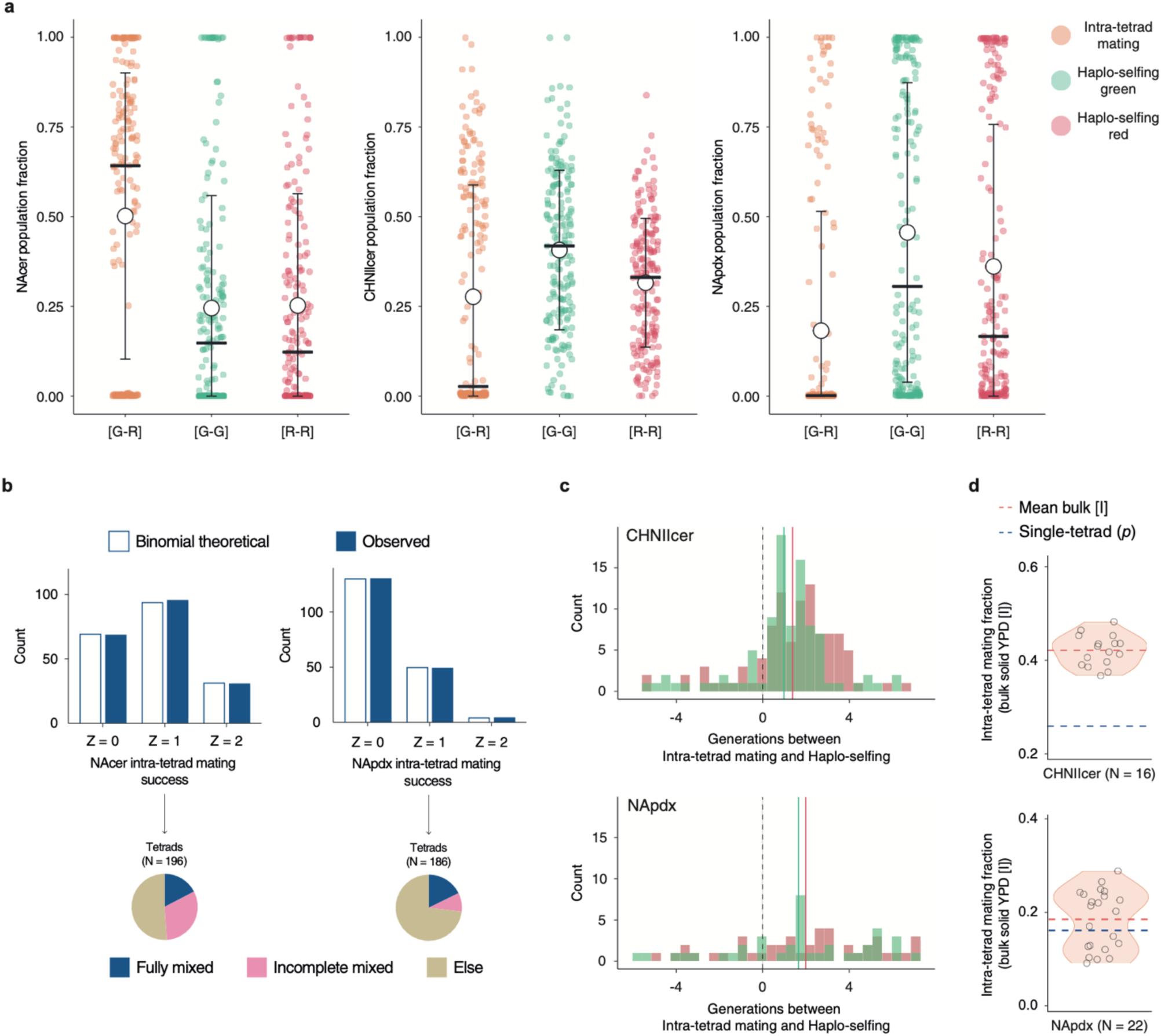
Tetrad-level heterogeneity in mating outcomes across diverse genetic backgrounds. **a**, Distribution of mating outcome across tetrads. Fractions of intra-tetrad mating [G–R], green haplo-selfed [G–G] and red haplo-selfed [R–R] derived populations are shown for each genetic background. Each point represents an individual tetrad (NAcer, *N* = 196; CHNIIcer, *N* = 191; NApdx, *N* = 186). White circles denote population means, whereas horizontal bars indicate medians. Differences between mean and median illustrate the asymmetric distributions of tetrad compositions, highlighting that tetrad-to-tetrad heterogeneity may be masked by population level metrics. **b**, Comparison between the observed distributions of intra-tetrad mating events per tetrad (Z) and the theoretical expectation under binomial model for NAcer and NApdx backgrounds. Insets summarize tetrads classified as fully mixed (all three classes detected), incompletely mixed (one class absent), or single outcome tetrads (only one class detected). **c**, Inferred number of generations by which green [G–G] and red [R–R] haplo-selfing lagged behind intra-tetrad mating. Analysis was restricted to tetrads with *Z* = 1, which contain descendants originating from both reproductive modes (CHNIIcer, *N* = 97; NApdx, *N* = 50; **Methods**). The vertical dashed line corresponds to synchronous mating events, whereas colored vertical lines indicate the mean inferred delay. **d**, Relationship between the probability of intra-tetrad mating estimated from the single-tetrad assay (*p*) and the corresponding intra-tetrad mating fraction ([I]) measured in bulk solid YPD-1X experiments for both CHNIIcer and NApdx genetic backgrounds.

## Methods

### Yeast strains and growth media

Strains used in this work are listed in Supplementary Table 1 (**Table S1**). The North American *S. cerevisiae* (YPS128, here NAcer) was used as the focal genetic background throughout the entire study. To benchmark gaMATe and compare mating outcomes across lineages, we additionally included a *S. cerevisiae* isolate from China (SX3, here CHNIIcer), and a *S. paradoxus* from North America (YPS138, here NApdx)^15,26^. All three backgrounds are wild, homozygous diploid isolates, except at the *MAT* locus. Unless otherwise stated, for each experiment, cells were revived from glycerol cryostocks on YPD agar (1% yeast extract, 2% peptone, 2% dextrose, 2% agar), and incubated overnight at 30°C.

### Genetic engineering

Strain engineering was performed following a standard lithium acetate/polyethylene glycol (LiAc/PEG) transformation protocol combined with CRISPR/Cas9-mediated genome editing, as previously described^27^. Briefly, cells were grown overnight and diluted 1:20 into 5 mL of fresh YPD and incubated for 4 hours at 30°C with shaking at 220 r.p.m. Cells were washed once with sterile distilled water and once with 0.1 M lithium acetate (LiAc). After complete removal of the LiAc solution, 240 µL of 50% PEG 3350 (Sigma-Aldrich), 36 µL of 1 M LiAc, 50 µL of PCR-amplified donor DNA, and >500 ng of CRISPR/Cas9 plasmid were added sequentially. The transformation mixture was vortexed for 2 minutes, incubated for 30 minutes at 30°C, and heat-shocked for 30 minutes at 42°C. Cells were then collected by centrifugation, resuspended in 1 mL fresh YPD, allowed to recover for 2 hours at 30°C, and plated on selective medium corresponding to the drug-resistance marker carried by the CRISPR/Cas9 plasmid. Mutant clones were streaked to single colonies on YPD plates without drug selection to promote spontaneous loss of the CRISPR/Cas9 plasmid. Genome-edited clones were systematically validated by PCR or, for allele replacements, by Sanger sequencing. Oligonucleotides, CRISPR guide RNAs, and Sanger sequencing services were provided by Eurofins Genomics.

#### Engineering of gaMATe at rainbow locus

The gaMATe system relies on the constitutive expression of fluorescent reporters integrated in close proximity to the *MAT* locus (∼2 kb, **Figure 1C**), at a site hereafter referred to as the *rainbow* locus. The *rainbow* locus was selected for its tight genetic linkage to *MAT*, thereby minimizing meiotic recombination between the fluorescent reporter and the mating-type locus and preserving the association between fluorescent identity and mating type across meiosis. The *rainbow* locus spans a neutral intergenic region on chromosome III, where a 13 bp sequence located 137-149 bp downstream of the *BUD5* ORF in *S. cerevisiae*, or a 10 bp sequence located 178-187 bp downstream of the *BUD5* ORF in *S. paradoxus*, was replaced with the fluorescent reporter cassette. Fluorescent reporter cassettes were PCR-amplified directly from plasmids p2B-60 (blue; pTDH3-yetSapphire-tADH1), p2B-65 (green; pTDH3-yomVenus-tADH1), and p2B-66 (red; pTDH3-yomCherry-tADH1) using long primers Bd-27/Bd-28 (NAcer), Bd-79/Bd-28 (CHNIIcer), or Bd-77/Bd-78 (NApdx), flanked with 50 bp homology arms targeting the *rainbow* locus. Genomic integration of the fluorescent cassettes was promoted by CRISPR/Cas9-mediated cleavage, as previously described^62^. Briefly, 20 bp CRISPR target sequences within the *rainbow* locus (*S. cerevisiae*, GATTACATCTAATTTACGTC; *S. paradoxus*, GTACTTAATCAACTACCTAA), each adjacent to a PAM (NGG), were selected. The corresponding guide sequences were cloned by Gibson Assembly (NEBuilder HiFi DNA Assembly) into the BsaI-digested pL58-KanMX plasmid backbone^62^, yielding CRISPR/Cas9 plasmids ChVp-rainb1 (*S. cerevisiae*) and ChVp-rainb2 (*S. paradoxus*). Strains were subsequently transformed as described above using >500 ng of CRISPR/Cas9 plasmid together with 50 µL of PCR-amplified donor DNA. To generate strains carrying both green and red fluorescent reporters, the green and red PCR-amplified donor DNAs were co-transformed, and clones displaying both fluorescent signals were PCR-validated (**Table S12**).

#### HO gene deletion from gaMATe strains

The *HO* gene was deleted by disrupting its ORF with a hygromycin resistance cassette PCR-amplified from pL-57 using long primers Bd-17/Bd-18 harboring 50 bp homology arms targeting *HO*. Genomic integration at the *HO* locus was promoted by CRISPR/Cas9 using plasmid pL88-KanMX-2xHO, derived from pUDP004 and engineered to carry kanamycin resistance instead of nourseothricin resistance, as previously described in^62^. Transformations were performed as detailed above using at least 500 ng of pL88-KanMX-2xHO plasmid together with 50 µL of PCR-amplified donor DNA. Transformants were selected on agar plates supplemented with G418 and hygromycin, streaked to single colonies, and validated by PCR to confirm correct genomic integration and complete loss of *HO* from both homologous chromosomes (**Table S12**).

#### Engineering of GPA1 variants in Nacer

To generate the alternative *GPA1* W82R and S469I alleles, two distinct CRISPR guide RNAs were designed to overlap the respective mutation sites, such that incorporation of the donor DNA both introduced the desired nucleotide substitutions and disrupted subsequent CRISPR/Cas9 recognition. The guide RNAs targeting *GPA1* W82R (TACAGTATGCTCAAGTGATA) and GPA1 S469I (AAAAAAGTGGTATTATATGA) were cloned into the BsaI-digested pL58-KanMX plasmid backbone^62^ by Gibson Assembly, yielding CRISPR/Cas9 plasmids ChVp-W82_GPA1 and ChVp-S469_GPA1, respectively. Donor DNAs consisted of 120 bp synthetic oligonucleotides spanning the Cas9 cleavage site and carrying the desired nucleotide substitutions. For *GPA1* W82R, the PAM sequence associated with the guide RNA directly overlapped codon W82. In the donor DNA, the A at nucleotide position 244 of *GPA1* coding sequence was replaced with G, and the C at position 246 was replaced with T. Together, these substitutions converted tryptophan to arginine while simultaneously disrupting the PAM sequence. For *GPA1* S469I, the PAM sequence was located in the 3’ UTR, 17 bp downstream of the mutation of interest. In the donor DNA, the C at nucleotide position 1406 of *GPA1* coding sequence was replaced with A, resulting in the substitution of serine with isoleucine, while the C at position 3 of the 3’ UTR was replaced with T to disrupt the PAM sequence. Each donor oligonucleotide was synthesized together with its reverse-complementary oligonucleotide to generate a double-stranded donor DNA (Bd-81/Bd-82 and Bd-83/Bd-84 for the W82R and S469I variants, respectively). Complementary oligonucleotides were mixed at a final concentration of 10 µM each into 50 µL of distilled water, heated for 10 minutes at 95°C, allowed to cool to room temperature, and subsequently added to the transformation mixture. Transformations and mutant validation were performed as described above (**Table S12**).

### Screening of gRNA target sites across *S. cerevisiae*

The conservation of gaMATe CRISPR/Cas9 target site across *S. cerevisiae* was assessed using the genome assemblies belonging to the 1011 project^15^. The complete 20 bp guide RNA sequence adjacent to the 3 bp of PAM was searched against each genome using fasta36 (v.36.3.8h), requiring 100% identity. For each strain, the number of exact matches was recorded. Target-site universality was defined as the percentage of strains for which at least one exact match was detected in the corresponding genome assembly.

### Experimental quantification of yeast mating using gaMATe

To implement gaMATe, we used diploid strains carrying defined combinations of fluorescent markers, hereafter referred to as [“Blue-Blue”] and [“Green-Red”] genotypes (**Figure 1D; Figure S1A-B; Appendix 1**). The two strains were sporulated separately to generate fluorescently labeled tetrads, which were subsequently mixed in equal proportion (50:50) to establish the mating mix (**Appendix 1**). Independent experiments were initiated from fresh cultures established from frozen stocks (**Table S1**). Unless otherwise stated, within each experiment, biological replicate cultures were subsequently divided into two or three technical replicates, which were analyzed separately (**Table S2**).

Cells were sporulated for at least 72 hours following the protocol described in^27^. Briefly, isolates were precultured in liquid YPD and diluted 1:50 into 10 mL of pre-sporulation medium (YPA; 2% potassium acetate, 1% yeast extract, 2% peptone), followed by incubation for 48 hours at 30°C with shaking at 220 r.p.m. Pre-sporulated cells were then transferred to 2% liquid potassium acetate (KAc) medium, and incubated for 72 hours at 23°C with shaking at 220 r.p.m. Sporulation efficiency was >99% before mating mixtures were established. To generate each mating mixture, 500 µL of each ascus genotype were combined and centrifuged, and the resulting pellet was resuspended in rich medium to allow spore germination and mating, resulting in a final volume of 1 mL. Before mixing, each sporulated culture had an OD₆₀₀ of ∼0.3 after 1:10 dilution (corresponding to approximately 10⁷cells/mL in the non-diluted cultures). The resulting 1 mL mating mixture was defined as the *standard* (1X) density. Solid YPD at the standard 1X density was defined as the *standard* mating environment. Mating assays were performed at 30°C for NAcer and CHNIIcer and at 23°C for NApdx. To assess the effect of local physical structure of the mating environment, identical mating mixes were exposed to solid, static liquid or shaken liquid YPD. Aliquots of the initial 1 mL mating mixture were used across the three conditions. For the liquid conditions, two 450 µL aliquots were each diluted 1:10 into fresh medium and incubated either statically or with shaking. For the solid condition, 5 µL of the remaining 100 µL of mating mixture was spotted onto agar medium, corresponding to approximately 2 x 10^5^ cells distributed over an area of ≈40 mm² (≈5200 cells/mm²). Following germination and mating (overnight for NAcer and CHNIIcer; 2 overnights for NApdx, reflecting the slower germination and subsequent growth of NApdx), the composition of the resulting diploid populations was quantified by flow cytometry according to the experimental scheme and gating strategy described in **Figure S1-2**. For the solid condition, cells were collected by sweeping across the entire diameter of the spot to minimize spatial sampling bias, resuspended in 200 µL of medium, and subsequently diluted in PBS for flow-cytometric analysis. Samples were analyzed using CytoFLEX LX and CytoFLEX S flow cytometers (Beckman Coulter), with at least 10,000 events recorded per sample. The relative frequencies of intra-tetrad mating, haplo-selfing, and inter-tetrad mating were estimated from the proportions of the corresponding gated fluorescent populations. Green, red, and sapphire fluorescence were detected using the B525-FITC-A, Y610-mCherry-A, and V525-KrO-A channels, respectively.

To assess the effects of medium composition and cell density, mating assays were performed in three media: the standard YPD and two additional mating environments, SOE (1% sucrose, 0.5% fructose, 0.5% glucose, 0.1% yeast extract and 0.15% peptone)^42^ and GAL (1% yeast extract, 2% peptone, 2% galactose)^63^. Five relative ascus densities were tested: 1X (standard density), 2X, 5X, 10X, and 25X. Higher relative densities were generated from the same starting amount of sporulated culture by reducing the final resuspension volume from 1 mL of the standard 1X condition to 500 µL, 200 µL, 100 µL, and 40 µL for the 2X, 5X, 10X, and 25X conditions, respectively. All vs all combinations of the three media and five cell densities were tested, with the standard condition (YPD, 1X density) included as an internal reference in each experimental batch. The composition of the resulting diploid populations was quantified by flow cytometry as described above. For NApdx in GAL, the incubation time was further extended because of its slower germination dynamics observed in this medium.

To assess the effect of ascus integrity on mating outcomes (Figure 2B-C), sporulated cultures were collected, washed with water, and resuspended in 200 µL of dissection buffer (2 M sorbitol, 10 mM EDTA, and 100 mM sodium phosphate) supplemented with 10 µL of zymolyase (5 mg/mL). Cultures were incubated at 30°C for either 0, 3, or 6 h, after which samples were assayed under solid or shaking YPD conditions.

### Experimental estimation of recombination between the *MAT* and *rainbow* loci

To assess the association between mating type and fluorescence identity, recombination between the *MAT* and the *rainbow* locus was experimentally quantified. An *HO*-deleted NAcer diploid strain carrying the [“Green-Red”] genotype was sporulated, and individual asci were dissected by micromanipulation. Following colony growth on the dissection plates, the fluorescent phenotype of each spore-derived colony was determined using the PhenoBooth+ (Singer). Mating type was subsequently determined by colony PCR using short primers GL_141, GL_142, and GL_143 with MATa and MATα identified based on amplicon sizes of 544 bp and 404 bp, respectively (**Figure S2**). A total of 580 haploid spores were analyzed. Comparison of fluorescent phenotype and *MAT* genotype identified [MATa, Red] or [MATα, Green] as the predominant associations, observed in 577/580 spores, thus classified as non-recombinant genotypes. The reciprocal [MATa, Green] or [MATα, Red] associations were detected in three spores and classified as recombinant genotypes, corresponding to an observed recombinant frequency of 0.52%.

### Estimation of the recombination parameter

Using the binom.test function from the R stats package, we computed the Clopper-Pearson exact 95% confidence interval for the observed recombinant frequency^64^, yielding an interval of *r* ∈ [0.107% - 1.504%]. For subsequent analyses, the recombination parameter (*r*) was fixed at 1%, a value within the experimentally estimated confidence interval (**Appendix 1**).

### Flow-cytometry validation of ploidy following germination and mating

Ploidy following germination and mating was assessed by propidium iodide (PI) staining and flow-cytometry analysis of DNA content, based on the protocol described in^15^, with modifications. Strains were recovered from glycerol stocks in 200 µL of liquid YPD in 96-well plates and grown overnight at 30°C. Cells were washed once and fixed overnight in 70% ethanol. Fixed cells were washed once and diluted 1:10 into staining solution containing 10 µg/mL PI (15µM), 100 µg/mL RNase A, and 0.1% (v/v) TritonX. Samples were incubated for at least 2 hours at 37°C before flow-cytometric analysis. At least 10,000 events per sample were recorded, with PI fluorescence detected using the Y585-PE-A channel. A haploid NAcer strain and the parental diploid NAcer [B-B] strain were used to define the DNA-content regions corresponding to the haploid and diploid G1 peaks, respectively. To determine whether residual haploid cells could bias fluorescence-based quantification of mating outcomes, ploidy was assessed for a selected set of [B-B] germination and mating conditions after overnight incubation, following the mating procedure described above (*Experimental quantification of yeast mating using gaMATe*; **Table S6**). The proportion of events falling within the haploid G1 DNA-content region (G1_n) was used to quantify residual haploid cells. Representative DNA-content distributions are shown in **Figure S2B**, with quantification across the tested conditions reported in **Table S6**.

### Growth phenotyping

Strains were cultivated overnight to saturation in rich medium and subsequently diluted 1:100 or 1:1000 into fresh YPD. Growth was monitored at 30°C over 48 hours by measuring optical density every 20 minutes using an Agilent LogPhase 600 (LP600) plate reader. Growth-curve processing and parameter estimation were performed in R using the gcplyr package (**Table S11**)^65^. OD values were blank-corrected, and the specific growth traits were estimated using the model free calc_deriv() function (percapita = TRUE, trans_y = “log”) with a five-timepoint sliding window. The maximum specific growth rate (μmax) was defined as the maximum smoothed per-capita derivative among measurements with a blank-corrected OD ≥0.1, whereas growth yield was defined as the highest blank-corrected OD reached by each curve.

### Phylogenetic analysis of *GPA1* alleles across *Saccharomyces*

The phylogenetic tree shown in **Figure 2D** was inferred from whole-genome assemblies of a selected panel of strains downloaded from ScRAPdb database (https://www.evomicslab.org/db/ScRAPdb/download/)^66,67^. The analysis included *S. cerevisiae* strains CRL (Wine/European clade), AEG (Bioethanol clade), BFP (Mediterranean Oak clade), the SGD reference strain, AKN (YPS128; North American clade), and BAH (CHNII clade), together with the North American *S. paradoxus* strain YPS138 and *S. jurei* strain NCYC3962 as outgroups. Whole-genome assemblies were processed using SKA (v0.5.1)^68^, with ska build and ska align used to generate the alignment. The resulting alignment was converted to FASTA format using SeqKit (v2.12.0)^69^ and used for maximum-likelihood phylogenetic inference with IQ-TREE 3 (v3.1.3)^70^. The best-fitting substitution model was selected using ModelFinder (-m MFP)^71^, and branch support was assessed using 1000 ultrafast bootstrap replicates and 1000 SH-aLRT replicates (-B 1000 -alrt 1000).

To assess the conservation of *GPA1* variants across the *Saccharomyces* genus, *GPA1* coding sequences (CDSs) from representative reference genomes were compared. The strains analyzed were YPS138 (North American *S. paradoxus*), CBS432 (European *S. paradoxus*), NCYC3962 (*S. jurei*), IFO1815-hc (*S. mikatae*), CR85 (*S. kudriavzevii*), H6 (*S. arboriculus*), CBS7001 (*S. uvarum*), and CL450 (S*. eubayanus*). CDS sequences of *GPA1* were aligned using MUSCLE in MEGA11 software^72,73^, and the resulting alignment was used to infer a maximum-likelihood tree under the Kimura two-parameter model^74^. The corresponding peptide sequences were also aligned in MEGA11, and amino-acid states at positions 82 and 469 were extracted from the protein alignment. The amino-acid states corresponding to the major alleles in *S. cerevisiae* were conserved across the analyzed *Saccharomyces* species, consistent with these being ancestral, while the minor *S. cerevisiae* alleles representing the derived states.

*<u>Illumina read mapping and genotyping</u>*: the distribution of *GPA1* alleles at amino-acid positions 82 and 469 was assessed across 3134 *S. cerevisiae* strains. Sequencing data were obtained from the ENA archive using iseq (v1.9.1)^75^ and combined with sequencing data from the 1011 Yeast Genomes collection^15^. Reads were aligned to the *S. cerevisiae* reference genome (SGD R64.3.1-20210421) using BWA-MEM (v0.7.17)^76^. Alignment files were processed by fixing mate information, marking duplicates, and coordinate sorting using SAMtools (v1.20)^77^, followed by variant calling using BCFtools (v1.20)^78^. Mapping and genotyping scripts were developed in the context of the fifth *Saccharomyces* Genome Resequencing Project (SGRP5). Individual gVCFs were merged, and the genomic positions corresponding to the codons encoding *GPA1* residues 82 and 469 were extracted. Extracted alleles were reverse-complemented using a custom R script before conversion of the resulting sequences to FASTA format using vcf2phylip.py (v2.9)^79^. Strains for which both positions could be unambiguously genotyped were retained, resulting in 3126/3134 for downstream analysis. Amino-acid allele combinations at positions 82 and 469 were analyzed using custom R scripts available at https://github.com/nicolo-tellini/GPA1.

### Single-ascus flow-cytometry of mating outcome

Individual tetrads obtained by sporulation of the engineered [“Green-Red”] diploid strains described above were isolated by micromanipulation, germinated in YPD plates until colony formation, and grown under conditions preventing inter-tetrad encounters (30°C for NAcer and CHNIIcer and 23°C for NApdx). Each resulting colony was individually resuspended in 200 µL YPD, diluted 1:10 in PBS, and analyzed by flow cytometry using the same instrument and fluorescence channels described above, with at least 10,000 events recorded per colony. Green-positive [G-G], Red-positive [R-R], and double-positive Green-Red [G-R] populations were quantified for each tetrad-derived colony as described in **Figure 4A**. Under the assay model, the single-color populations represent descendants arising from the two haplo-selfing outcomes, whereas the double-color positive population represents intra-tetrad mating products. For each ascus-derived colony, the frequencies of the three populations were normalized to sum to 1 and used for subsequent analyses (**Table S7**).

*<u>Determination of the number of intra-tetrad events (Z)</u>:* for each ascus, the number of intra-tetrad mating events was denoted as Z = 0, 1, or 2. Tetrads were classified as Z = 0 when the double-positive [G-R] population represented <1% of events (mating outcome corresponded to exclusive haplo-selfing); as Z = 2 when the combined frequency of the two single-positive populations ([G-G] + [R-R]) represented <1% of events (mating outcome corresponded to exclusive intra-tetrad mating); and as Z = 1 otherwise (mixed mating outcome was observed). To test whether the two compatible spore pairs within a tetrad behaved independently and had the same probability of undergoing intra-tetrad mating, the observed distribution of Z was compared, separately for each genetic background, with a binomial model. Specifically, Z was assumed to follow a Binomial(2, *p*) distribution, where *p* represents the probability that a compatible spore pair undergoes intra-tetrad mating. For each genetic background, *p* was estimated from the observed data as:

- *p* = [0 x n_obs(Z = 0) + 1 x n_obs(Z = 1) + 2 x n_obs(Z = 2)] / 2*N*,

where *N* corresponds to the total number of tetrads analyzed, 2*N* is the total number of compatible spore pairs, and n_obs(Z = 0), n_obs(Z = 1), and n_obs(Z = 2) are the observed numbers of tetrads classified as Z = 0, Z = 1, and Z = 2, respectively.

Given *p*, the expected numbers of tetrads in each category under the binomial model were calculated as:

- n_th(Z = 0) = *N* x (1 − *p*)^2^,
- n_th(Z = 1) = *N* x 2 x *p* x (1 − *p*), and
- n_th(Z = 2) = *N* x *p*^2^.

Finally, observed and expected numbers of tetrads in the three Z categories were compared using a Chi-square goodness-of-fit test with one degree of freedom.

*<u>Estimation of relative lineage-expansion between intra-tetrad mating and haplo-selfing (Δt_[G-R],[G-G]_ or Δt_[G-R],[R-R]_)</u>*: in tetrads classified as Z = 1, the delay, expressed in mitotic generations, between the initiation of the proliferation of the intra-tetrad-derived lineage and the expansion of the haplo-selfing-derived ones was estimated using the final relative abundances of the mating outcomes (**Appendix 3**). Assuming complete spore viability, these tetrads comprise three independently expanding mating-derived lineages: one [G-R] lineage resulting from intra-tetrad mating and two single-color lineages, [G-G] and [R-R], resulting from haplo-selfing. Single-color lineages representing <0.5% of the final population were excluded from the corresponding lineage-specific timing comparison; thus, when only one single-color lineage exceeded this threshold, the tetrad was retained and the timing difference was estimated using only the detectable lineage. Assuming equivalent growth rates among the three lineages after vegetative growth initiation (**Figure S1E**), the difference in the number of generations of expansion between two lineages can be inferred from their final abundance ratio. For a lineage originating from a single founder cell and doubling once per generation, its population size after *t* generations is given by n = 2^t^, and therefore t = log_2_(*n*). Consequently, the difference in expansion onset times between two lineages, (*a*) and (*b*), can be inferred directly from their final population sizes *n_a_* and *n_b_*: Δ*t*_(a,b)_ = *t*_a_ - *t*_b_ = log_2_(*n_a_*) − log_2_(*n_b_*) = log_2_(*n_a_*/*n_b_*). Because the ratio *n_a_*/*n_b_* is unchanged when absolute cell numbers are replaced by their corresponding population fractions, Δ*t*_(a,b)_ could be calculated directly from the relative abundances measured by flow cytometry. Accordingly, the relative difference in expansion between the intra-tetrad [G-R] lineage compared with each detectable haplo-selfed lineage was estimated as *Δt_[G-R],[G-G]_ or Δt_[G-R],[R-R]_* (**Appendix 3**). Positive values indicate that the [G-R] lineage underwent more generations of expansion before the common sampling time and, under the equal-growth-rate assumption, initiated expansion earlier than the corresponding haplo-selfed lineage. For this analysis, lineage expansion was defined from the beginning of vegetative proliferation rather than strictly from the fusion event. The [G-R] lineage was considered to originate from the zygote generated by intra-tetrad mating, whereas for each haplo-selfed lineage, the onset of expansion was defined by the first budding of the haploid spore. This framework assumes comparable haploid and diploid mitotic-cycle durations and that cell fusion occurs on a timescale short relative to a mitotic cycle.

### Spore viability

To quantify spore viability, 200 µL of sporulated cultures were collected, washed in water and resuspended in 200 µL of dissection buffer (sorbitol 2 M, EDTA 10 mM, sodium phosphate 100 mM), supplemented with 10 µL of zymolyase 5 mg/mL, and incubated 2 hours at 30°C. A total of 95, 96, and 72 tetrads were dissected for NAcer, CHNIIcer, and NApdx, respectively, on YPD agar plates using a SporePlay+ dissection microscope (Singer) (**Table S8**). Plates were incubated at 30°C for at least 48 hours, and spore viability was calculated as the proportion of dissected spores that formed visible colonies.

### Agar pad time-lapse microscopy

Tetrad germination and mating were monitored by time-lapse microscopy on agar pads. Agar pads were prepared as previously described in^80^: briefly, molten medium was placed into the wells of a custom 3D-printed mold positioned on a 50 mm round coverslip (Epredia, CB005005A140MNZ0). A second coverslip was immediately placed on top of the mold to flatten the pad surfaces and ensure a uniform optical path. After 15 minutes of solidification at room temperature, the bottom coverslip was removed and 4 µL of sporulated culture was deposited onto the surface of each pad. After the droplets had dried, a new coverslip was placed on the mold, and the assembled device was mounted on the microscope. Timelapse imaging initiated approximately 1 hour after tetrads were deposited on the pads, accounting for device preparation and selection of imaging positions. Images were acquired at a 5-minutes interval for 12 hours at 30°C, using the NIS-Elements software on a Nikon Eclipse Ti2 inverted microscope equipped with a Hamamatsu ORCA-Flash4.0LT Digital camera and a Nikon Plan Apo Lambda 60X/1.40 Oil objective. Acquisition settings were as follow: phase contrast, 50 ms, 50% intensity; mVenus (470 nm excitation & 519 nm emission filters), 200 ms, 50% intensity; mCherry (555 nm excitation & 630 nm emission filters), 200 ms, 50% intensity.

*<u>Image analysis</u>*: time-lapse microscopy data were manually annotated at single-tetrad resolution. Asci distributed across multiple imaging positions were analyzed for each genetic background (*N* = 62 for NAcer, *N* = 53 for CHNIIcer, and *N* = 63 for NApdx). For each ascus, gamete release was scored as the first time point showing visible rupture of the ascus wall and release of the enclosed gametes. The time of the first mating event was scored as the first time point showing irreversible cell fusion between the corresponding gametes, and the delay between gamete release and the first mating event was calculated for each tetrad as Δt = t_first mating − t_gamete release. Temporal measurements were expressed relative to the placement of tetrads on the agar pads, accounting for the ∼1 hour interval between sample deposition and the onset of image acquisition. For each spore, mating outcomes were additionally classified as intra-tetrad mating or haplo-selfing. Cell fusion was identified primarily based on characteristic zygote morphology; intra-tetrad mating events were additionally supported by the emergence of the double-positive [G-R] fluorescence phenotype resulting from cytoplasmic mixing between the green and red mating partners. The mating outcome of the first and second mating events was recorded, together with any observed gamete-death event. Representative time-lapse sequences are provided in **Video S1-2**. For visualization of the supplementary videos, lookup tables (LUTs) were adjusted relative to the corresponding time-lapse images to improve contrast and brightness; these display adjustments were applied for visualization only and were not used for event annotation or quantitative analysis.

### Experimental synchronization of germination timing

Hybridization between CHNIIcer and NApdx was quantified under asynchronous and experimentally synchronized germination conditions by selectively advancing germination of NApdx, the slower-germinating background identified by single-tetrad time-lapse microscopy (**Figure 5B**). Two pre-germination intervals, 4.5 and 7 hours, were selected from the distribution of NApdx gametes release times: the first one approximated the mean of the distribution, whereas the second corresponded to one of the latest observed time points. Sporulated NApdx cultures were centrifuged, resuspended in YPD at the standard 1X condition described above, and incubated statically at 23°C for 4.5 or 7 hours. A 0-hour treatment served as the asynchronous control, whereas throughout the pre-germination interval, sporulated CHNIIcer cultures remained under the original pre-assay conditions. At the indicated time point, 500 µL of each culture was separately centrifuged, and each pellet was resuspended in 500 µL of the medium corresponding to the experimental condition. The two cultures were then combined to a final volume of 1 mL, corresponding to the standard 1X density. Mating was assayed on solid medium across the same medium-density conditions described above. CHNIIcer and NApdx carried the [B-B] and [G-R] genotypes, respectively. Hybrid diploids were identified by flow cytometry using the logic and the gating strategy described in **Figure S1A-B**, which discriminated intra-species hybrids from the alternative diploid mating outcomes (e.g., inter-tetrad events within the same background).

### Quantification and statistical analysis

Data processing, visualization, and statistical analyses were performed using R v2021.09.2 and GraphPad Prism v11.0.2. Definitions of *N* and summary statistics are reported in the corresponding figure legend or accompanying text. Unless otherwise indicated, statistical significance was defined as *p-value* < 0.05. To quantify differences in mating strategy composition among experimental conditions, the frequencies of the mating outcomes were averaged across replicates to obtain a mean compositional vector for each condition (**Table S3**). Pairwise Jensen–Shannon divergence was calculated between the mean compositional vectors using the JSD() function from the R package philentropy v0.9.0. Jensen–Shannon distance (JSdist) was subsequently calculated as the square root of the divergence. Pairwise JSdist values are visualized as distance matrices with and without hierarchical clustering (**Figure S4**). Flow-cytometry data were analyzed using CytExpert v2.4.0.28. Figures were prepared using BioRender and Adobe Illustrator, and genetic constructs were designed using Geneious v9.1.8.

## Lead contact

Requests for further information and resources should be directed to and will be fulfilled by the lead contact, Gianni Liti

## Data availability

Data supporting the findings of this study are available within the article or the supporting information. Any additional raw data or additional information required to reanalyze the data are available upon request. Plasmids will be deposited on Addgene and will become freely available upon publication for academic research only (**Table S13**). All the yeast strains are available on request (**Table S1**).

## Acknowledgment

We thank Christian Braendle, Anthony Long, Jason Isheng Tsai, and Jonas Warringer for critical reading of the manuscript. This work was supported by the Agence Nationale de la Recherche (ANR-20-CE12-0020 to G.L., G.F., B.L., and M.C.L.; ANR-18-CE12-0004, ANR-20-CE13-0010, ANR-23-CE12-0031, ANR-24-CE12-7740, and ANR-15-IDEX-01 to G.L.); the Fondation Bettencourt Schueller (Impulscience to G.L.); the Fondation pour la Recherche Médicale (EQU202003010413 to G.L.); SATT Sud-Est (to G.L.); the Associazione Italiana per la Ricerca sul Cancro (AIRC Investigator Grant IG 2024, ID 30391 to M.C.L.); the Ministero dell’Università e della Ricerca (PRIN 2022, ID P20224N9X4 to M.C.L.); the Université Côte d’Azur IDEX (to B.P.B.); and the Swiss National Science Foundation (SNSF) through the NCCR Microbiomes Fellowship (to C.V.), the NCCR Microbiomes Grant 51NF40_180575 (to S.M. and J.S.L.), and the Eccellenza Grant 181272 (to C.VU.). The authors acknowledge IRCAN’s Cytometry Core Facility (CYTOMED), supported by le Cancéropole PACA, la Région PACA, le Conseil Départemental 06, INSERM, ARC, IBiSA, and the Conseil Départemental 06 de la Région PACA.

## Declaration of generative AI and AI-assisted technologies in the manuscript preparation process

during the preparation of this work, the authors used ChatGPT (OpenAI) to check grammar and to improve the readability of the text. The authors reviewed and edited the content and take full responsibility for the final manuscript.

## Author contributions

Conceptualization, C.V., B.L., M.C.L., G.F., G.L.; Methodology, C.V., B.P.B., N.V., S.P., C.VU., J.S.L., M.C.L; Formal Analysis, C.V., N.V., M.D.C., S.P., N.T., S.K.; Investigation, C.V., with input from B.P.B., K.T., S.L.; Resources, B.P.B., S.M., B.L., M.C.L., G.F., G.L.; funding acquisition, B.L., M.C.L., G.F., G.L.; Writing - original draft C.V., G.L. All authors reviewed the manuscript and approved the final version for submission.

## Supplementary Information

### Supplementary Appendices

<u>Appendix 1:</u> Formal description of the mathematical framework used to quantify the relative contributions of mating strategies within a population.

<u>Appendix 2:</u> Mathematical modelling incorporating selection and model fitting to experimental data.

<u>Appendix 3:</u> Mathematical framework for the single-tetrad flow-cytometry assay

### Supplementary time-lapse movies

<u>Video S1-2:</u> representative time-lapse microscopy of NAcer tetrads germinating and mating on YPD agar pads.

### Supplementary Tables

*All Supplementary Tables are available for download*:

<u>Table S1:</u> *Saccharomyces* strains used in this study.

<u>Table S2:</u> Complete flow-cytometry dataset, including raw percentages and inferred fractions of intra-tetrad mating, haplo-selfing, and inter-tetrad mating.

<u>Table S3:</u> Pairwise Jensen-Shannon distances.

<u>Table S4:</u> *GPA1* aa82/aa469 combinations across *S. cerevisiae*.

<u>Table S5:</u> Mating outcome in *ΔHO* experimental subset.

<u>Table S6:</u> Assessment of haploid cells upon germination and mating.

<u>Table S7:</u> Normalized flow-cytometry measurements from the single-tetrad assay.

<u>Table S8:</u> Gamete viability measured by asci micromanipulation.

<u>Table S9:</u> Dataset underlying image analysis.

<u>Table S10:</u> Flow-cytometry dataset for the targeted synchronization experiment.

<u>Table S11:</u> Growth parameters.

<u>Table S12:</u> Oligonucleotides used in work.

<u>Table S13</u>: Plasmids used in this work.

