## Appendix 1 for "A reproductive hierarchy controls mating fates among sibling yeast gametes"

#### Overview of the mathematical framework

We developed a mathematical framework to (i) quantify the relative contributions of different mating strategies within a population and (ii) to characterize how these vary across genotypes and environmental conditions. Specifically, this method maps experimentally measured proportions of cells carrying different color combinations to the frequencies of intra-tetrad mating, haplo-selfing, and inter-tetrad mating, enabling direct interpretation as percentages within the population. This mapping accounts for both the recombination rate and the asymmetries arising from color combinations that can result from multiple mating outcomes (**Figure S1-2**).

#### Framework and notations

Square brackets [“.”] denote the genotype of a cell. For haploid cells, the genotype is specified at the mating type (*MAT*) locus (MATa or MAT$\alpha$) and at the *rainbow* locus (**Figure 1c**), where fluorescent markers are expressed (Blue = mT-Sapphire, Red = mCherry, Green = mVenus). For diploid cells, the genotype at the *MAT* locus is omitted, as it is assumed to be MATa/MAT$\alpha$.

To generate gametes, [”Blue-Blue”] and [”Green-Red”] diploids are induced to sporulate separately. We assume that all cells enter meiosis and form tetrads (denoted by curly bracket {.}). The ancestral population is constructed by mixing a proportion $q \in ]0,1[$ of tetrads derived from the sporulated [”Blue-Blue”] population with a proportion $1-q$ of tetrads from the sporulated [”Green-Red”] population. The resulting asci mixture is transferred onto a rich medium, allowing spores to germinate and undergo mating.

A haploid cell can mate with a cell of the opposite mating-type to form a diploid cell. The mating partner may descend from (i) a spore belonging to an unrelated tetrad (inter-tetrad mating or outcrossing), (ii) another spore within the same tetrad (intra-tetrad mating), or (iii) the same spore following mating-type switching (haplo-selfing). We assume that mating-type switching occurs only in haploid gametes and leads to haplo-selfing.

In our framework, both haploid and diploid cells can divide mitotically, and descendants of individual spores are not constrained to follow identical trajectories. We further assume that the allele at the *rainbow* locus has no effect on cellular fitness (**Figure S1**). In addition, the growth rate of the diploid cells is homogeneous, being independent of both the allele at the *rainbow* locus and the mating process that produces the first diploid of each lineage (**Figure S1**). Typically, this assumption implies that diploids [“Blue-Blue”] and diploids [“Green-Red”] are genetically identical and homozygous, except at the *rainbow* and the *MAT* loci.

We assume that the population at the time of measurement consists exclusively of diploid cells (**Figure S2**).

#### Recombination between the rainbow locus and the MAT locus

In diploid cells from the pre-meiotic [“Green-Red”] population, the Red allele at the *rainbow* locus is linked to the MATa allele, whereas the alternative Green allele is linked to the MAT$\alpha$ allele. During meiosis, recombination between the *rainbow* locus and the *MAT* locus may occur with probability $r \in ]0, 0.5[$ (**Figure S2**). As a result, haploid products can carry either parental non-recombinant ({[MATa, Red] or [MAT$\alpha$, Green]}) or recombinant ({[MATa, Green] or [MAT$\alpha$, Red]}) combinations of alleles, with respective probabilities of $1 - r$ and $r$. Neglecting multiple crossover events between the *rainbow* and *MAT* loci within a single meiosis, tetrads can be classified into two categories:

- non-recombinant tetrads ($\bar{\rho}$), in which all spores retain parental linkage configurations {[MATa, Red], [MATa, Red], [MAT$\alpha$, Green, [MAT$\alpha$, Green]}, in proportion $1 - 2r$;
- recombinant tetrads ($\rho$), in which two spores carry recombinant combinations {[MATa, Green], [MAT$\alpha$, Red], [MATa, Red], [MAT$\alpha$, Green]}, in proportion $2r \in ]0, 1[$.

The value of the recombination parameter $r$ has been estimated experimentally (**Figure S2**).

### Inference of mating strategy contributions

#### Composition of the ancestral population

We first formalize the composition of the ancestral population which defines the distribution of genotypes entering the mating process. Tetrads are sampled uniformly at random from this population. We denote by $T_{\mathrm{BB}}$, $T_{\mathrm{RG},\bar{\rho}}$ and $T_{\mathrm{RG},\rho}$ the events that a sampled tetrad corresponds to a {[MATa, Blue], [MATa, Blue], [MAT$\alpha$, Blue], [MAT$\alpha$, Blue]} tetrad, a non-recombinant {[MATa, Red], [MATa, Red], [MAT$\alpha$, Green], [MAT$\alpha$, Green]} tetrad, and a recombinant {[MATa, Green], [MAT$\alpha$, Red], [MATa, Red], [MAT$\alpha$, Green]} tetrad respectively. The associated probabilities depend on both the mixing proportion $q$ and the recombination parameter $r$, and are given by:

$$\mathbb{P}(T_{\mathrm{BB}}) = q, \qquad \mathbb{P}(T_{\mathrm{RG},\bar{\rho}}) = (1 - 2r)(1 - q), \qquad \mathbb{P}(T_{\mathrm{RG},\rho}) = 2r(1 - q).$$

We next consider sampling at the level of individual gametes. Let $G_s$ denote the event that a randomly sampled spore carries genotype ["$G_s$"], spanning the set of possible haploid combinations of alleles at the *MAT* and *rainbow* loci.

$$G_s \in \{[\mathrm{MATa}, \mathrm{Blue}], [\mathrm{MAT}\alpha, \mathrm{Blue}], [\mathrm{MATa}, \mathrm{Red}], [\mathrm{MAT}\alpha, \mathrm{Green}], [\mathrm{MAT}\alpha, \mathrm{Red}], [\mathrm{MATa}, \mathrm{Green}]\}$$

The probabilities associated with these events are:

$$\mathbb{P}([\mathrm{MATa}, \mathrm{Blue}]) = \mathbb{P}([\mathrm{MAT}\alpha, \mathrm{Blue}]) = \frac{\mathbb{P}(T_{\mathrm{BB}})}{2} = \frac{q}{2},$$

$$\mathbb{P}([\mathrm{MATa}, \mathrm{Red}]) = \mathbb{P}([\mathrm{MAT}\alpha, \mathrm{Green}]) = \frac{\mathbb{P}(T_{\mathrm{RG},\bar{\rho}})}{2} + \frac{\mathbb{P}(T_{\mathrm{RG},\rho})}{4} = \frac{(1 - r)(1 - q)}{2},$$

$$\mathbb{P}([\mathrm{MAT}\alpha, \mathrm{Red}]) = \mathbb{P}([\mathrm{MATa}, \mathrm{Green}]) = \frac{\mathbb{P}(T_{\mathrm{RG},\rho})}{4} = \frac{r(1 - q)}{2}.$$

#### Composition of the final population

We next characterize the composition of the final population, corresponding to the observable distribution of fluorescent phenotypes after mating. Cells are sampled uniformly at random from this population. Let $G$ denote the event that a sampled cell carries genotype [“$G$”] where $G$ is defined by the combination of alleles at the *MAT* and *rainbow* loci.

$$G \in \{[\text{Blue-Blue}], [\text{Blue-Red}], [\text{Blue-Green}], [\text{Red-Green}], [\text{Red-Red}], [\text{Green-Green}]\}$$

We denote by $\mathbb{P}(G)$ the corresponding associated probabilities. Each diploid cell in the final population originates from a single founding mating event occurring at the onset of its lineage. We distinguish three possible mating strategies: inter-tetrad mating (outcrossing, event $O$), intra-tetrad mating (event $I$), and haplo-selfing (event $H$). These occur with probabilities $\mathbb{P}(O)$, $\mathbb{P}(I)$, and $\mathbb{P}(H)$, respectively, satisfying

$$\mathbb{P}(O) + \mathbb{P}(I) + \mathbb{P}(H) = 1$$

The distribution of genotypes in the final population can therefore be expressed using the law of total probability as:

$$\forall G, \quad \mathbb{P}(G) = \mathbb{P}(G \cap H) + \mathbb{P}(G \cap I) + \mathbb{P}(G \cap O)$$

and the chain rule as:

$$\forall G, \quad \mathbb{P}(G) = \mathbb{P}(H)\mathbb{P}_H(G) + \mathbb{P}(I)\mathbb{P}_I(G) + \mathbb{P}(O)\mathbb{P}_O(G)$$

This formulation establishes a direct link between the observable genotype distribution and the underlying contributions of each mating strategy.

#### Conditional probabilities linking mating outcomes to observed phenotypes

We next derive the conditional probabilities that link each mating events to the resulting genotype and fluorescent phenotypes in the final population (= color combinations, **Figure S1**). We first consider haplo-selfing, in which a haploid cell undergoes mating-type switching and subsequently mates with itself to form a diploid cell. Under this mechanism, both copies of the diploid genome originate from the same haploid progenitor. As a result, only homozygous single-color genotypes can be generated, and genotypes combining different fluorescent markers are not observed.

$$\mathbb{P}_H([\text{Blue-Red}]) = \mathbb{P}_H([\text{Blue-Green}]) = \mathbb{P}_H([\text{Red-Green}]) = 0.$$

For a given single-color diploid, this probability ($\mathbb{P}_H$) is equal to the expected frequency of an ancestral haploid carrying the corresponding allele, which is given by the frequency of that allele among haploid spores in the initial population:

$$\mathbb{P}_H([\text{Blue-Blue}]) = \mathbb{P}([\text{MATa}, \text{Blue}]) + \mathbb{P}([\text{MAT}\alpha, \text{Blue}]) = q,$$
$$\mathbb{P}_H([\text{Red-Red}]) = \mathbb{P}([\text{MATa}, \text{Red}]) + \mathbb{P}([\text{MAT}\alpha, \text{Red}]) = 1 - q,$$
$$\mathbb{P}_H([\text{Green-Green}]) = \mathbb{P}([\text{MATa}, \text{Green}]) + \mathbb{P}([\text{MAT}\alpha, \text{Green}]) = 1 - q.$$

We next consider intra-tetrad mating, in which two haploid spores originating from the same tetrad fuse to form a diploid cell. $\mathbb{P}_I(G)$ is the probability that a cell has the genotype ["$G$"] if the first diploid cell in its ancestry was formed by intra-tetrad mating. Using the law of total probabilities and the chain rule:

$$\mathbb{P}_I(G) = \sum_{T \in \{T_{\text{BB}}, T_{\text{RG},\bar{\rho}}, T_{\text{RG},\rho}\}} \mathbb{P}(T)\mathbb{P}_{I\cap T}(G),$$

where $\mathbb{P}_{I\cap T}(G)$ is the probability to have genotype $G$ from intra-tetrad inbreeding in a tetrad $T$. Therefore, we obtain:

$$\mathbb{P}_I([\text{Blue-Red}]) = \mathbb{P}_I([\text{Blue-Green}]) = 0,$$
$$\mathbb{P}_I([\text{Blue-Blue}]) = \mathbb{P}(T_{\text{BB}}) = q,$$
$$\mathbb{P}_I([\text{Red-Green}]) = \mathbb{P}(T_{\text{RG},\bar{\rho}}) \times 1 + \mathbb{P}(T_{\text{RG},\rho}) \times \frac{1}{2} = (1-q)(1-r),$$
$$\mathbb{P}_I([\text{Red-Red}]) = \mathbb{P}_I([\text{Green-Green}]) = \mathbb{P}(T_{\text{RG},\rho}) \times \frac{1}{4} = \frac{1}{2}r(1-q).$$

Finally, we consider inter-tetrad mating, in which two haploid cells originating from distinct tetrads fuse to form a diploid cell. $\mathbb{P}_O(G)$ is the probability that a cell has genotype ["$G$"] if the first diploid in its ancestry was formed by outcrossing. This depends on the probability to sample compatible spores of each color as:

$$\mathbb{P}_O([\text{Blue-Red}]) = \mathbb{P}([\text{MATa}, \text{Blue}])\mathbb{P}([\text{MAT}\alpha, \text{Red}]) + \mathbb{P}([\text{MATa}, \text{Red}])\mathbb{P}([\text{MAT}\alpha, \text{Blue}]) = q(1-q),$$

$$\mathbb{P}_O([\text{Blue-Green}]) = \mathbb{P}([\text{MATa}, \text{Blue}])\mathbb{P}([\text{MAT}\alpha, \text{Green}]) + \mathbb{P}([\text{MATa}, \text{Green}])\mathbb{P}([\text{MAT}\alpha, \text{Blue}]) = q(1-q),$$

$$\mathbb{P}_O([\text{Red-Green}]) = \mathbb{P}([\text{MATa}, \text{Green}])\mathbb{P}([\text{MAT}\alpha, \text{Red}]) + \mathbb{P}([\text{MATa}, \text{Red}])\mathbb{P}([\text{MAT}\alpha, \text{Green}]) = ((1-q)^2((1-r)^2 + r^2),$$

$$\mathbb{P}_O([\text{Blue-Blue}]) = \mathbb{P}([\text{MATa}, \text{Blue}])\mathbb{P}([\text{MAT}\alpha, \text{Blue}]) = q^2,$$

$$\mathbb{P}_O([\text{Red-Red}]) = \mathbb{P}([\text{MATa}, \text{Red}])\mathbb{P}([\text{MAT}\alpha, \text{Red}]) = (1-q)^2 r(1-r),$$

$$\mathbb{P}_O([\text{Green-Green}]) = \mathbb{P}([\text{MATa}, \text{Green}])\mathbb{P}([\text{MAT}\alpha, \text{Green}]) = (1-q)^2 r(1-r).$$

##### Estimates for $q$, $\mathbb{P}(H)$, $\mathbb{P}(I)$ and $\mathbb{P}(O)$

: Using the observed frequencies of fluorescence combinations in the final population, we compute the estimates for $q$, $\mathbb{P}(H)$, $\mathbb{P}(I)$ and $\mathbb{P}(O)$. These relationships are formalized as a system of equations that relates the observed frequencies as functions of the underlying contribution of each mating strategy:

$$\forall G, \quad \mathbb{P}(G) = \mathbb{P}(H)\mathbb{P}_H(G) + \mathbb{P}(I)\mathbb{P}_I(G) + \mathbb{P}(O)\mathbb{P}_O(G)$$

$$\Longrightarrow \begin{cases} \mathbb{P}([\text{Green-Green}]) & = \frac{1-q}{2}\mathbb{P}(H) + \frac{(1-q)r}{2}\mathbb{P}(I) + (1-q)^2 r(1-r)\mathbb{P}(O) \\ \mathbb{P}([\text{Red-Red}]) & = \frac{1-q}{2}\mathbb{P}(H) + \frac{(1-q)r}{2}\mathbb{P}(I) + (1-q)^2 r(1-r)\mathbb{P}(O) \\ \mathbb{P}([\text{Blue-Blue}]) & = q\mathbb{P}(H) + q\mathbb{P}(I) + q^2\mathbb{P}(O) \\ \mathbb{P}([\text{Red-Green}]) & = (1-q)(1-r)\mathbb{P}(I) + (1-q)^2((1-r)^2 + r^2)\mathbb{P}(O) \\ \mathbb{P}([\text{Blue-Green}]) & = q(1-q)\mathbb{P}(O) \\ \mathbb{P}([\text{Blue-Red}]) & = q(1-q)\mathbb{P}(O) \end{cases}$$

$$\Longrightarrow \begin{cases} \frac{\mathbb{P}([\text{Green-Green}]) + \mathbb{P}([\text{Red-Red}])}{2} & = \frac{1-q}{2}\mathbb{P}(H) + \frac{(1-q)r}{2}\mathbb{P}(I) + (1-q)^2 r(1-r)\mathbb{P}(O) \\ \mathbb{P}([\text{Blue-Blue}]) & = q\mathbb{P}(H) + q\mathbb{P}(I) + q^2\mathbb{P}(O) \\ \mathbb{P}([\text{Red-Green}]) & = (1-q)(1-r)\mathbb{P}(I) + (1-q)^2((1-r)^2 + r^2)\mathbb{P}(O) \\ \frac{\mathbb{P}([\text{Blue-Green}]) + \mathbb{P}([\text{Blue-Red}])}{2} & = q(1-q)\mathbb{P}(O) \end{cases}$$

$$\Longrightarrow \begin{cases} q & = \mathbb{P}(\text{Blue-Blue}]) + \frac{\mathbb{P}([\text{Blue-Green}]) + \mathbb{P}([\text{Blue-Red}])}{2} \\ \mathbb{P}(H) & = \frac{\mathbb{P}([\text{Blue-Blue}])}{q} - \frac{\mathbb{P}([\text{Red-Green}])}{(1-q)(1-r)} \\ & \quad + \frac{(1-q)((1-r)^2 + r^2) - q(1-r)}{q(1-q)(1-r)} \frac{\mathbb{P}([\text{Blue-Green}]) + \mathbb{P}([\text{Blue-Red}])}{2} \\ \mathbb{P}(I) & = \frac{\mathbb{P}([\text{Red-Green}])}{(1-q)(1-r)} + \frac{-((1-r)^2 + r^2)}{q(1-r)} \frac{\mathbb{P}([\text{Blue-Green}]) + \mathbb{P}([\text{Blue-Red}])}{2} \\ \mathbb{P}(O) & = \frac{1}{q(1-q)} \frac{\mathbb{P}([\text{Blue-Green}]) + \mathbb{P}([\text{Blue-Red}])}{2} \end{cases}$$

#### Stability of the estimates following diploidization

We finally verify that the genotype probabilities inferred from the framework remain invariant during subsequent population growth. Specifically, we show that once all cells in the population are diploid, the probabilities $\mathbb{P}(H)$, $\mathbb{P}(I)$ and $\mathbb{P}(O)$ are constant over time under exponential or logistic (*e.g.*, with saturation) growth dynamics.

Let $t_0 \in \mathbb{R}^+$ denote the first time point at which all cells in the population are diploid. We denote by $n(t)$ the total population size at $t$, and by $n_G(t)$ the number of cells of genotype $G$ at time $t$. We consider two growth regimes

##### Exponential growth

We first assume that the population grows exponentially. Under the assumption that all diploid cells share the same growth rate $c \in \mathbb{R}^+$, the population dynamics satisfies:

$$\forall t \geq t_0, \quad n(t) = n(t_0)e^{c(t-t_0)} \quad \text{and} \quad n_G(t) = n_G(t_0)e^{c(t-t_0)}$$

It follows that, the frequency of the population that carries genotype $G$ at time $t$ is:

$$\forall t \geq t_0, \quad f_G(t) = \frac{n_G(t)}{n(t)} = f_G(t) = \frac{n_G(t_0)e^{c(t-t_0)}}{n(t_0)e^{c(t-t_0)}} = \frac{n_G(t_0)}{n(t_0)}$$

##### Saturated growth

We next consider logistic growth, leading to saturation at carrying capacity $K$. Assuming that all diploid cells share the same intrinsic growth rate $c \in \mathbb{R}^+$, the population dynamics satisfies:

$$\forall t \geq t_0, \begin{cases} n'(t) & = cn(t)(1 - \frac{n(t)}{K}) \\ n'_G(t) & = cn_G(t)(1 - \frac{n(t)}{K}) \end{cases}$$

We denote as $f_G(t) = \frac{n_G(t)}{n(t)}$ the fraction of the population that carries genotype $G$ et time $t$.

$$\forall t \geq t_0, \qquad f'_G(t) = \frac{n'_G(t)n(t) - n_G(t)n'(t)}{n^2(t)} = 0$$

$$\implies \forall t \geq t_0, \qquad f_G(t) = f_G(t_0) = \frac{n_G(t_0)}{n(t_0)}$$

Together, these results show that once all mating events have occurred and the population is fully diploid ($t \geq t_0$), genotype frequencies are invariant under both growth regimes. Consequently, the estimates for $\mathbb{P}(H)$, $\mathbb{P}(I)$ and $\mathbb{P}(O)$ derived from the observed population are also constant and independent of the time at which measurements are performed.
