## Appendix 2 for "A reproductive hierarchy controls mating fates among sibling yeast gametes"

### Appendix 2 for Vischioni *et al.*: Hierarchical mating-allocation models

This appendix describes the mathematical models used for the analysis presented in "Environmental context and population density shape mating outcomes through hierarchical allocation of reproductive modes", concerning the hierarchical mating-allocation.

#### Mating routes and their possible organization

The model considers three mating routes, **intra-tetrad mating**, **haplo-selfing**, and **inter-tetrad mating**, and experiments carried out at different spore densities and for different environments and strains. At tetrad density $x$, the frequencies of different routes are denoted by $P_{\text{intra}}$, $P_{\text{haplo}}$, and $P_{\text{inter}}$, respectively, with

$$P_{\text{intra}}(x) + P_{\text{haplo}}(x) + P_{\text{inter}}(x) = 1. \tag{1}$$

As density increases, the encounter probability of tetrades increases, and inter-tetrad mating becomes more common. The key question is therefore which of the other two routes gives way. If one route remains approximately stable while another decreases, the mating routes are not equivalent but follow an ordered, or hierarchical, organization. To investigate this, we considered four mathematical models embodying different scenarios. Each of these models makes a simple prediction about what should remain unchanged as density increases.

In the simplest reference case, which provides a null model for the absence of a hierarchical structure, the relative frequencies of all three mating routes are independent of tetrad concentration. In other words, changing the density of tetrads does not change the balance between intra-tetrad mating, haplo-selfing, and inter-tetrad mating.

Instead, a hierarchical organization introduces an order among the different mating routes. In the models, the route that has priority is assumed to occur with a probability that does not depend on tetrad concentration. The remaining two routes then compete for the fraction of mating events that is left after the preferred route has been accounted for.

The following equations define the models.

- **Null model: Control**
  All mating probabilities are independent of tetrad density $x$.

$$P_{\text{intra}} = p, \qquad P_{\text{inter}} = A, \qquad P_{\text{haplo}} = 1 - A - p. \tag{2}$$

- **Model 1: Intra-tetrad mating first**
  The probability of intra-tetrad mating, $P_{\text{intra}}$, is independent of tetrad density $x$. The sum of inter-tetrad and haplo-selfing probabilities is therefore constrained by normalization, while $P_{\text{inter}}$ varies with $x$.

$$P_{\text{intra}} = p, \qquad P_{\text{inter}} = f(x), \qquad P_{\text{haplo}} = 1 - f(x) - p. \tag{3}$$

- **Model 2: Haplo-selfing mating first**
  The probability of haplo-selfing, $P_{\text{haplo}}$, is independent of tetrad density $x$. The sum of intra-tetrad and inter-tetrad probabilities is therefore constrained by normalization, while $P_{\text{inter}}$ varies with $x$.

$$P_{\text{haplo}} = p, \qquad P_{\text{inter}} = f(x), \qquad P_{\text{intra}} = 1 - f(x) - p. \tag{4}$$

- **Model 3: Inter-tetrad mating first**
  The probability of inter-tetrad mating, $P_{\text{inter}}$, varies with tetrad density $x$. The remaining probability, $1 - P_{\text{inter}}$, is partitioned between intra-tetrad mating and haplo-selfing with a constant ratio.

$$P_{\text{inter}} = f(x), \qquad P_{\text{intra}} = \big(1 - f(x)\big)\, p, \qquad P_{\text{haplo}} = \big(1 - f(x)\big)(1 - p). \tag{5}$$

Here, $f(x)$ denotes a generic function, which depends on encounter probability, describing the dependence of the inter-tetrad mating probability on tetrad density. Its functional form is not specified at this stage, but a physical interpretation is given below.

#### A simple test of the mating hierarchy

Even without knowing how $f(x)$ depends on tetrad density, we can identify which model best explains the data using simple summary plots. The four models can be compared by analyzing the changes in intra-tetrad, inter-tetrad, and haplo-selfing probabilities relative to the reference density $x = 1$, defined as

$$\Delta P_{\text{intra}}(x) = P_{\text{intra}}(x) - P_{\text{intra}}(1), \quad (6)$$

$$\Delta P_{\text{haplo}}(x) = P_{\text{haplo}}(x) - P_{\text{haplo}}(1), \quad (7)$$

$$\Delta P_{\text{inter}}(x) = P_{\text{inter}}(x) - P_{\text{inter}}(1). \quad (8)$$

None of these relationships depends on the form of $f(x)$, hence the predictions are general. The tables below give the trajectory each model predicts in two diagnostic plots. The first diagnostic uses intra-tetrad mating, $\Delta P_{\text{intra}}$ vs. $\Delta P_{\text{inter}}$.

| **Model** | **Trajectory in** $(\Delta P_{\text{inter}}, \Delta P_{\text{intra}})$ | **Interpretation** |
|---|---|---|
| 1: Intra-tetrad first | $(\Delta P_{\text{inter}},\ 0)$ | Horizontal line: $P_{\text{intra}}$ unchanged |
| 2: Haplo-selfing first | $(\Delta P_{\text{inter}},\ -\Delta P_{\text{inter}})$ | Line through the origin with slope $-1$ |
| 3: Inter-tetrad first | $(\Delta P_{\text{inter}},\ -p\,\Delta P_{\text{inter}})$ | Line through the origin with slope $-p$ |
| Null model (control) | $(0,\ 0)$ | A single point at the origin |

Table 1: Predicted behavior with $\Delta P_{\text{inter}}$ on the horizontal axis and $\Delta P_{\text{intra}}$ on the vertical axis. These “preference trajectories” are independent of the form of $f(x)$, which encodes encounter probability.

The second diagnostic uses haplo-selfing in place of intra-tetrad mating, so we plot $\Delta P_{\text{haplo}}$ against $\Delta P_{\text{inter}}$. Again, each model predicts a specific preference trajectory, which does not depend on $f(x)$.

| **Model** | **Trajectory in** $(\Delta P_{\text{inter}}, \Delta P_{\text{haplo}})$ | **Interpretation** |
|---|---|---|
| 1: Intra-tetrad first | $(\Delta P_{\text{inter}},\ -\Delta P_{\text{inter}})$ | Line through the origin with slope $-1$ |
| 2: Haplo-selfing first | $(\Delta P_{\text{inter}},\ 0)$ | Horizontal line: $P_{\text{haplo}}$ unchanged |
| 3: Inter-tetrad first | $(\Delta P_{\text{inter}},\ -(1-p)\,\Delta P_{\text{inter}})$ | Line through the origin with slope $-(1-p)$ |
| Null model (control) | $(0,\ 0)$ | A single point at the origin |

Table 2: Predicted trajectories with $\Delta P_{\text{inter}}$ on the horizontal axis and $\Delta P_{\text{haplo}}$ on the vertical axis. As in the first diagnostic, the preference trajectories do not depend on the form of $f(x)$.

For each condition, we can estimate $p$, which defines the slopes in the diagnostic plots for Model 3, by pooling the measured fractions across the densities tested:

$$\hat{p} = \frac{\sum_x \bar{P}_{\text{intra}}(x)}{\sum_x \bar{P}_{\text{intra}}(x) + \sum_x \bar{P}_{\text{haplo}}(x)}, \quad (9)$$

where $\bar{P}$ denotes the replicate mean. To visualize these predictions, we plot the change in each mating fraction relative to the reference density $x = 1$ against the corresponding change in inter-tetrad mating (Figure 1; same representation as main-text Figure 3C). Each model produces a characteristic pattern.

The data clearly depart from the null model. As density increases, inter-tetrad mating rises and the other routes change systematically in all three genetic backgrounds, with the exception of NApdx on solid GAL, where inter-tetrad mating remains close to 1% at all densities. Among the hierarchical alternatives, the overall pattern is closest to Model 1: haplo-selfing decreases as inter-tetrad mating increases, whereas the intra-tetrad fraction does not decrease and, in YPD, tends to increase. The quantities predicted to remain constant under Models 2 and 3 instead show stronger systematic variation.

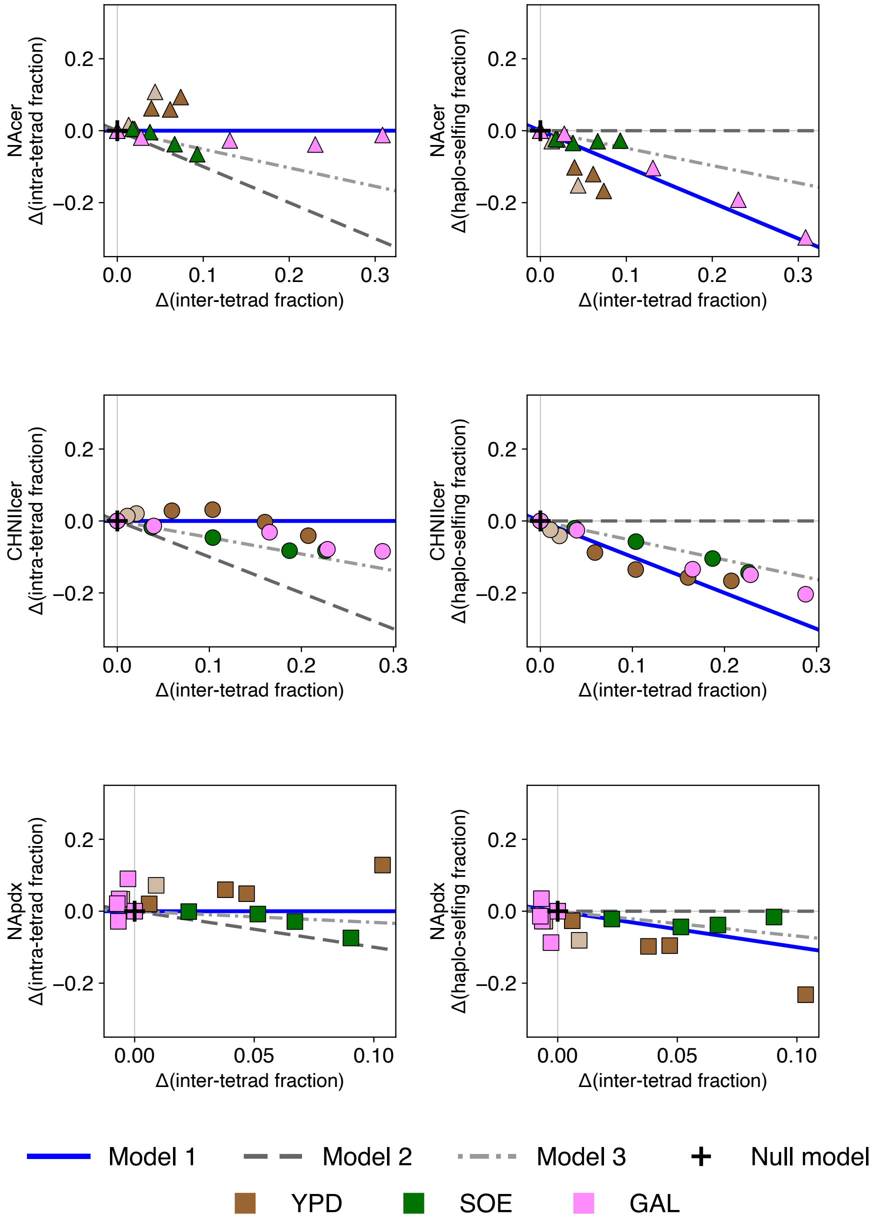


Figure 1: **Changes in the intra-tetrad fraction (left column) and in the haplo-selfing fraction (right column) plotted against the change in inter-tetrad mating, relative to the reference density** $x = 1$. Rows correspond to the strains NAcer, CHNIIcer and NApdx. Model 1 predicts a flat relationship in the left column, in which intra-tetrad mating is retained as density increases. Model 2, , in which haplo-selfing is retained, predicts a flat relationship in the right column. Model 3 predicts that the two routes decrease together in a fixed proportion, with slopes $-\hat{p}$ and $-(1-\hat{p})$, respectively. Here $\hat{p}$ is estimated directly from the measured fractions as described in the text and averaged over the four conditions of each background. The null model predicts no change and therefore corresponds to a single point at the origin.

#### Testing of the mating hierarchy in each condition

To test each condition separately, we chose a simple empirical function based on the observed behavior. Inter-tetrad mating is rare at low density, increases with density, and eventually approaches a plateau:

$$P_{\text{inter}}(x) = P_{\max}\left(1 - e^{-\beta x}\right). \tag{10}$$

Here, $P_{\max}$ determines the maximum fraction of mating events that can be inter-tetrad ($A$ in Table 3), while $\beta$ determines how rapidly this level is approached as density increases.

On solid media, this functional form has a simple physical interpretation. Spores deposited on agar are immobile, so two tetrads can mate only if they lie within an effective contact distance $r$, set by the spore spacing, the extent of microcolony growth before spores commit to intra-tetrad mating or haplo-selfing, and the range of pheromone signalling. If tetrads are deposited at random with surface density $\rho \propto x$, the number of neighbours within distance $r$ is Poisson distributed, and the probability that a tetrad has at least one potential partner is

$$\phi(x) = 1 - e^{-\pi r^2 \rho} = 1 - e^{-\beta x}, \qquad \beta = \pi r^2 \rho_1, \tag{11}$$

where $\rho_1$ is the surface density at the reference concentration $x = 1$. Encounters are therefore negligible when the typical distance between tetrads is much larger than $r$, where $\phi(x) \simeq \beta x$, and saturate once this distance becomes comparable to $r$. In this interpretation, $\beta$ measures the contact area relative to the area available per tetrad at $x = 1$, and $P_{\max}$ is the fraction of mating events that are inter-tetrad among tetrads that do have a neighbour. The fitted values of $\beta$ ($\approx$ 0.2-0.5 on solid media) place the crossover between these two regimes within the tested density range. By contrast, in shaking liquid cultures inter-tetrad mating changes by at most a few percent over the available range of densities (Figures 3-5), suggesting that encounters are rare under these conditions.

For each dataset, we compared the measured fractions of the three mating routes, averaged over replicates, with the model predictions at every density tested. We quantified the mismatch using the relative entropy (Kullback–Leibler divergence),

$$D(x) = \sum_m \bar{P}_m(x) \ln\left[\frac{\bar{P}_m(x)}{P_m(x)}\right], \tag{12}$$

where $m$ indexes the three mating routes. The fitting score was

$$L = -N \sum_x D(x), \tag{13}$$

with $N = 10^4$, the average number of cells processed in the FACS analysis in each condition. The best-fitting parameters were identified by an exhaustive search over the ranges given in Table 3. The three hierarchical models have three fitted parameters, $p$, $A$, and $\beta$, whereas the null model has two. Because models with more parameters can generally fit data more closely, we compared them using the Bayesian information criterion (BIC),

$$\text{BIC} = k \ln(3n) - 2L, \tag{14}$$

where $k$ is the number of fitted parameters and $n$ is the number of density values tested. The factor $3n$ corresponds to the three mating-route frequencies measured at each density. For each dataset, BIC values were converted into relative weights,

$$w_i = \frac{\exp\left[-(\text{BIC}_i - \text{BIC}_{\min})/2\right]}{\sum_j \exp\left[-(\text{BIC}_j - \text{BIC}_{\min})/2\right]}. \tag{15}$$

The condition-specific analysis gives a more nuanced picture (Figure 2; individual fits in Figures 3–5). The preferred model varies across conditions, with Model 1 providing the most frequent description, while Model 3 is preferred in a subset of conditions. Model 2 is not preferred in any dataset, and the null model is preferred in only one. Thus, while the overall pattern is consistent with the model-independent analysis, the preferred hierarchical organization is not identical across all environments.

| Model | $p$ range | $A$ range | $\beta$ range |
|---|---|---|---|
| 1: Intra-tetrad first | 0.01 to 0.99 (step 0.01) | 0.01 to $1-p$ (step 0.01) | 0 to 10 (step 0.1) |
| 2: Haplo-selfing first | 0.01 to 0.99 (step 0.01) | 0.01 to $1-p$ (step 0.01) | 0.01 to 5.01 (step 0.1) |
| 3: Inter-tetrad first | 0.01 to 0.99 (step 0.01) | 0.01 to 0.99 (step 0.01) | 0 to 10 (step 0.1) |
| Null model | 0.01 to 0.99 (step 0.01) | 0.01 to $1-p$ (step 0.01) | – (not used) |

Table 3: Parameter ranges explored for each model in the condition specific model testing

The spatial picture above also clarifies why the preferred model alternates between Model 1 and Model 3 across solid-media conditions. If only tetrads with a neighbour can engage in inter-tetrad mating, the population is a mixture of isolated tetrads (fraction $1-\phi$) and tetrads in contact (fraction $\phi$), and each observed frequency takes the form

$$P_m(x) = \big(1-\phi(x)\big)\,P_m^{\mathrm{iso}} + \phi(x)\,P_m^{\mathrm{c}}, \qquad m \in \{\mathrm{intra}, \mathrm{haplo}, \mathrm{inter}\}, \tag{16}$$

with $P_{\mathrm{inter}}^{\mathrm{iso}} = 0$. Since all frequencies are linear in $\phi(x)$, any allocation rule among tetrads in contact produces a straight line through the origin in the diagnostic plots of Figure 1, with slope $s = \Delta P_{\mathrm{intra}}/\Delta P_{\mathrm{inter}} = (P_{\mathrm{intra}}^{\mathrm{c}} - P_{\mathrm{intra}}^{\mathrm{iso}})/P_{\mathrm{inter}}^{\mathrm{c}}$, i.e. the amount of intra-tetrad mating lost per inter-tetrad mating event gained. Models 1, 2 and 3 correspond to the particular values $s = 0$, $s = -1$ and $s = -p$, respectively. The empirical slopes on solid media (least-squares fits through the origin) lie between these reference values and are distributed rather continuously: $-0.05$ (CHNIIcer, YPD), $-0.10$ (NAcer, GAL), $-0.29$ (CHNIIcer, GAL), $-0.40$ (CHNIIcer, SOE), $-0.57$ (NApdx, SOE) and $-0.58$ (NAcer, SOE), compared with $-\hat{p}$ between $-0.23$ and $-0.71$. Model selection among the three discrete hierarchies therefore assigns each condition to the closest reference slope, and conditions with similar behaviour (e.g. CHNIIcer and NAcer on GAL) can end up being attributed to different models. We can therefore interpret the hierarchical models as reference cases of a continuum: on solid media, the data are consistent with intra-tetrad mating being largely protected ($-p \le s \le 0$), with haplo-selfing always contributing most of the frequency recruited by inter-tetrad mating.

Importantly, this condition-specific comparison relies on the additional assumption that the density dependence of inter-tetrad mating can be described by the function $P_{\mathrm{inter}}(x) = P_{\max}\left(1-e^{-\beta x}\right)$. All four models share a further structural assumption, namely that density affects mating allocation only through the encounter probability $f(x)$, i.e. that the balance between intra-tetrad mating and haplo-selfing changes only in response to changes in inter-tetrad mating, so that intra-tetrad mating either remains constant or decreases as inter-tetrad mating increases. Several datasets suggest that this assumption does not always hold. In four datasets, NAcer and NApdx in YPD (both shaking and solid), the measured intra-tetrad fraction instead increases with density, by an amount larger than the increase in inter-tetrad mating (empirical slopes $s$ between $+1.2$ and $+4.5$), so that haplo-selfing declines in favour of both other routes. In shaking liquid this shift occurs while inter-tetrad mating changes by only a few percent, showing that it is not mediated by encounters between tetrads. These observations suggest a second, encounter-independent effect of density on the choice between intra-tetrad mating and haplo-selfing, for instance through germination timing, medium conditioning or density-dependent signalling. The preference for Model 1 in these datasets should therefore be read as the closest description within a class of models that does not include this additional effect. In NApdx on solid GAL, inter-tetrad mating shows no density dependence. In this case, the fitted parameters lie at the edges of the explored ranges, and the preference for the null model reflects the absence of a density response. Since the same strain shows a clear density response on solid YPD and SOE, we interpret this result as suggesting that in this condition haplo-selfing accounts for most of mating events at all densities and may pre-empt mating with neighbouring tetrads (so that the effective contact range $r$ in Eq. (11), and hence $\beta$, vanishes).

Taken together, these analyses show that mating routes are not used in fixed proportions as population density changes. The dominant pattern is that haplo-selfing is the route that gives way as density increases, while intra-tetrad mating is retained or, in YPD, increases. The condition-specific analysis supports the idea that the degree to which intra-tetrad mating is protected from inter-tetrad mating varies across environments, ranging from full protection (Model 1) to proportional competition (Model 3).

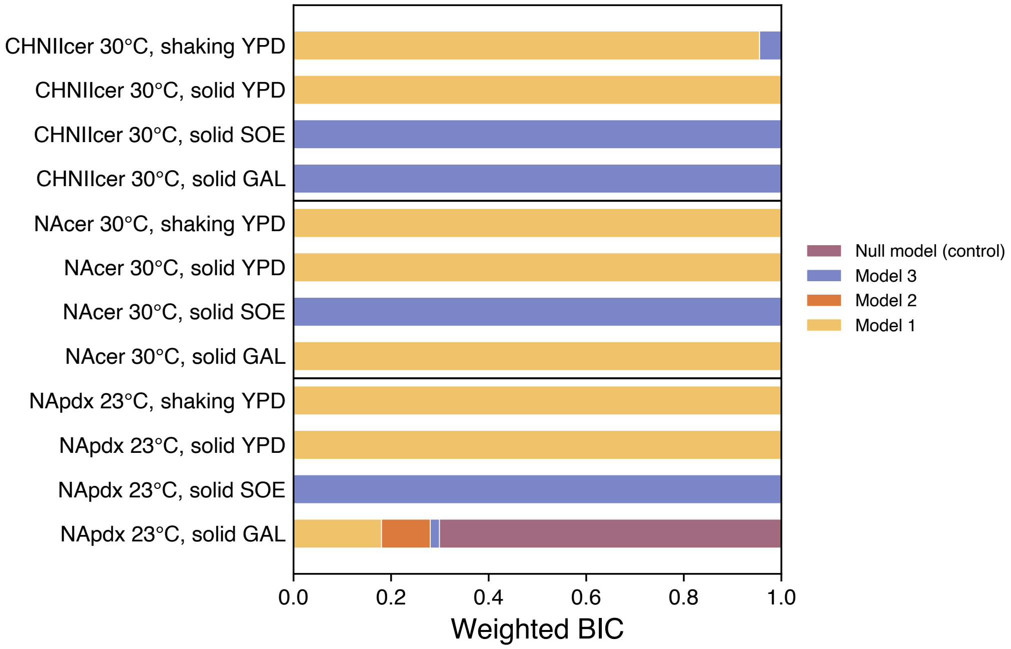


Figure 2: Relative support for the four models in each of the 12 datasets, based on BIC weights. Model 1 (intra-tetrad first) is preferred in seven of the 12 datasets, four of which show an increase in intra-tetrad mating that no model captures (see text), while Model 3 (inter-tetrad first) is preferred in a subset of conditions. Model 2 (haplo-selfing first) is not preferred in any dataset. The null model is preferred in only one dataset.

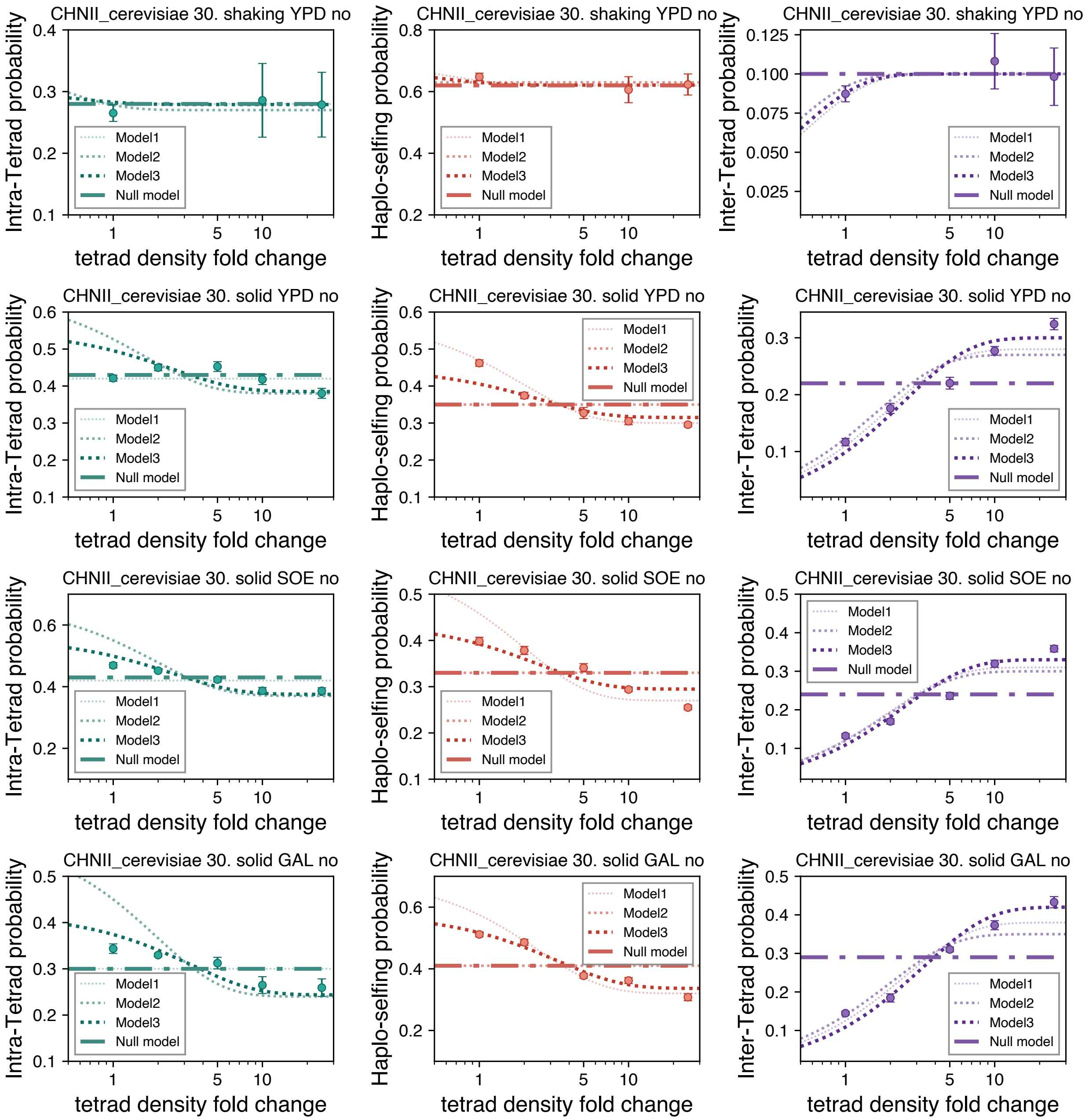


Figure 3: Model fits for the four *S. cerevisiae* CHNII conditions. Points show the mean ± s.e.m. across replicates at each density; lines show the fitted prediction of each model.

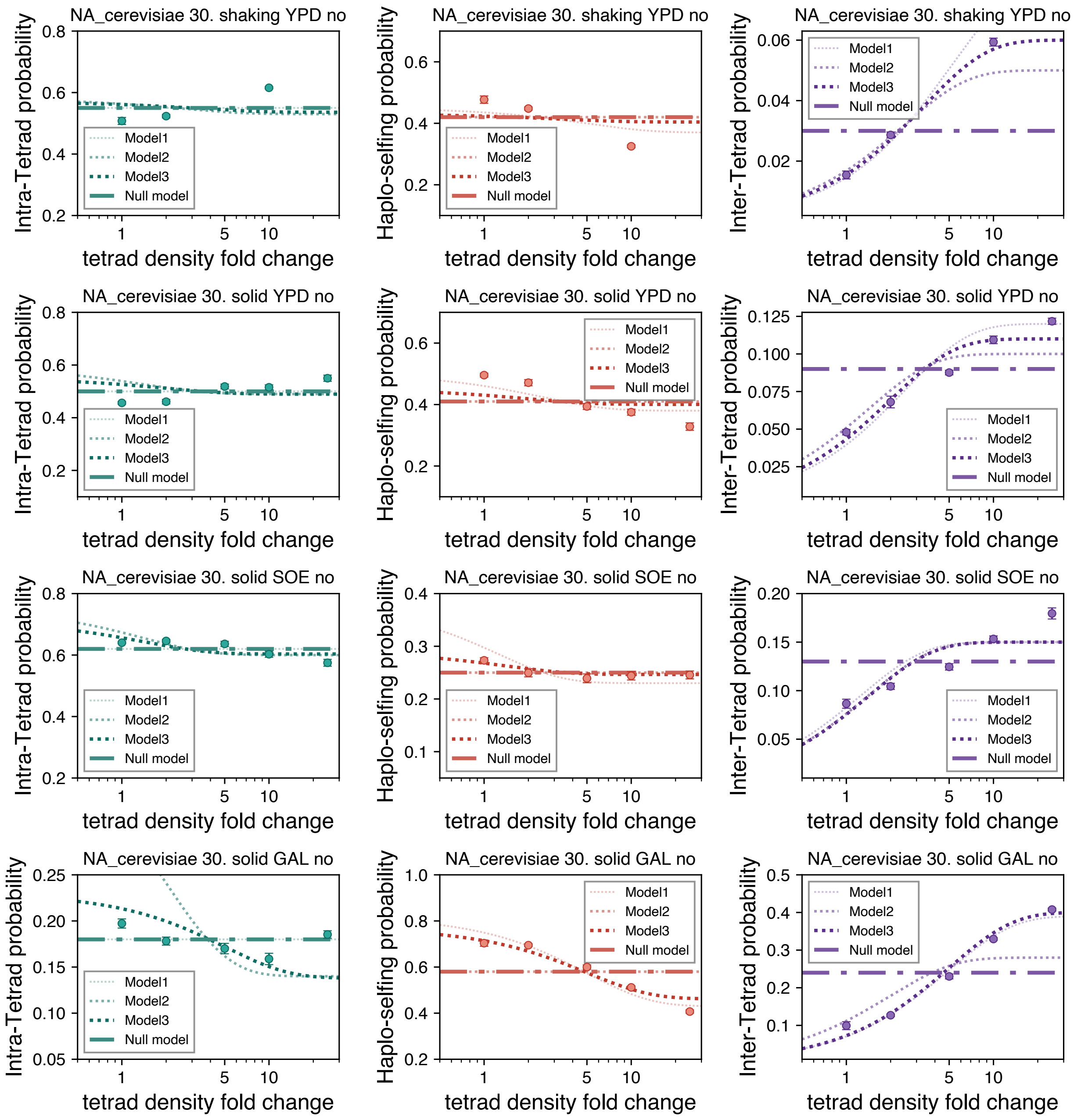


Figure 4: Model fits for the four *S. cerevisiae* NA conditions, with the same conventions as Figure 3.

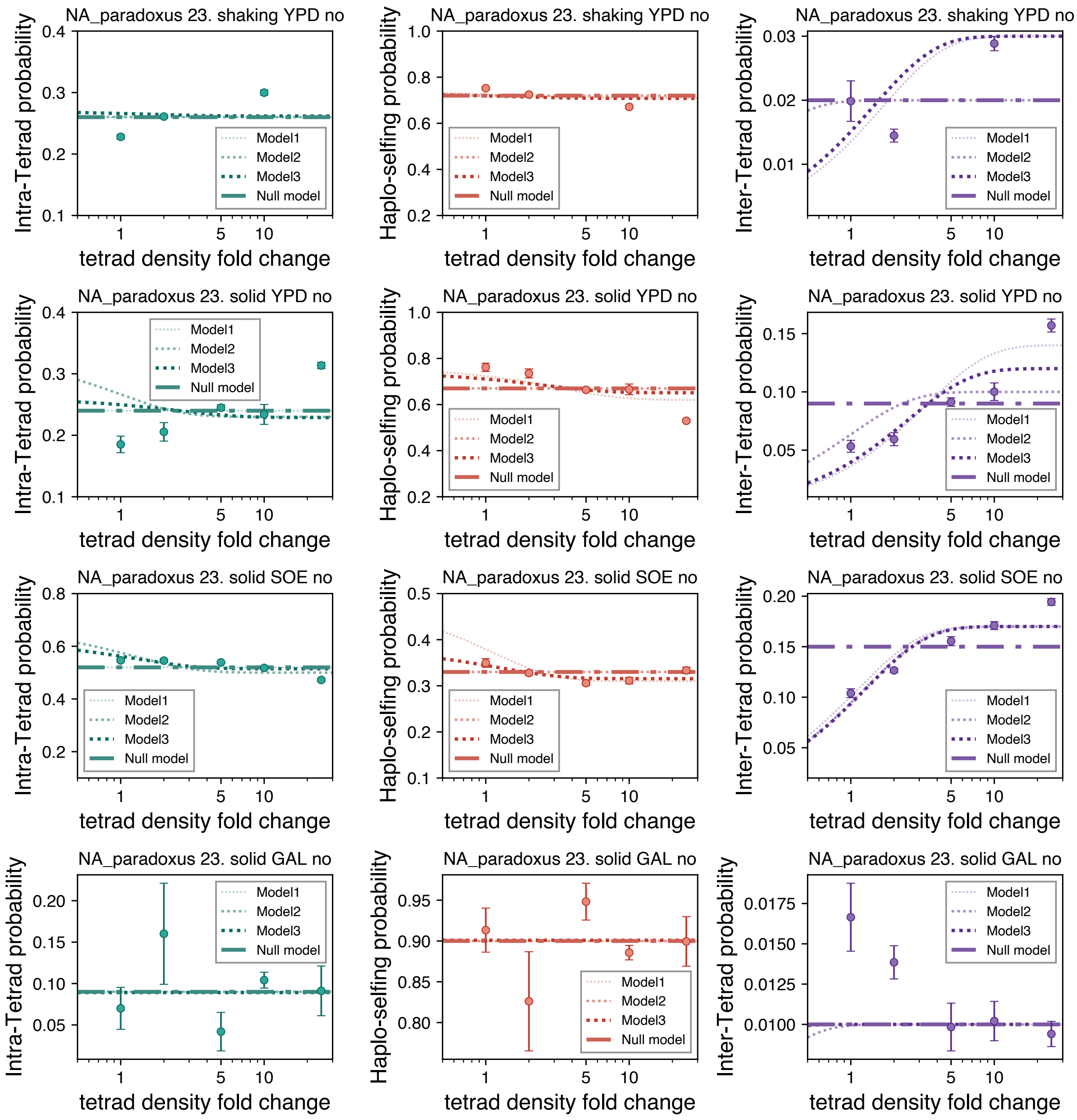


Figure 5: Model fits for the four *S. paradoxus* NA conditions, with the same conventions as Figure 3.
