## Appendix 3 for "A reproductive hierarchy controls mating fates among sibling yeast gametes"

#### Framework

A single tetrad containing two haploid red spores and two haploid green spores is isolated on YPD-agar, allowing spore germination, mating, and subsequent mitotic growth. The relative abundances of the fluorescent populations composing each tetrad-derived colony are quantified by flow-cytometry. We denote:

$N_{[R-R]}$ as the number of observed Red-Red cells ([R-R]), produced by haplo-selfing within a red-spore lineage followed by vegetative growth and expansion,

$N_{[G-G]}$ as the number of observed Green-Green cells ([G-G]), produced by haplo-selfing within a green-spore lineage followed by vegetative growth and expansion,

$N_{[G-R]}$ as the number of observed Green-Red cells ([G-R]), produced by intra-tetrad mating between a green spore and a red spore followed by vegetative growth and expansion.

We further denote

$p_{[R-R]} = \dfrac{N_{[R-R]}}{N_{\text{tot}}}$ as the proportion of [R-R] cells,

$p_{[G-G]} = \dfrac{N_{[G-G]}}{N_{\text{tot}}}$ as the proportion of [G-G] cells,

$p_{[G-R]} = \dfrac{N_{[G-R]}}{N_{\text{tot}}}$ as the proportion of [G-R] cells.

We assume that intra-tetrad mating occurs after germination and before any mitotic division. We therefore define $Z$ as the number of intra-tetrad mating events occurring within a tetrad, with

$$Z \in \{0, 1, 2\}.$$

If all four spores survive and contribute to the resulting population, we expect

tetrads with $Z = 0$ to have $p_{[G-R]} = 0$,

tetrads with $Z = 1$ to have $0 < p_{[G-R]} < 1$,

tetrads with $Z = 2$ to have $p_{[G-R]} = 1$.

In the following analysis, we focus on $Z = 1$ tetrads, defined in practice as tetrads for which

$$0.01 \leq p_{[G-R]} \leq 0.99.$$

These tetrads exhibit a mixed mating strategy: a fraction of the population originates from an intra-tetrad mating event between a green and a red spore, whereas the remaining lineages arise from haplo-selfing.

### Population Size Ratios, and Delays in Growth Initiation

#### General theory

Consider a population of cells growing vegetatively at a constant growth rate and doubling once at each generation. Starting from a single cell at generation 0, the population size after $t$ generations is

$$n = 2^t,$$

and therefore the number of generations since the initiation of vegetative growth is

$$t = \log_2(n).$$

Now, consider two populations, $A$ and $B$, growing at the same rate, with population sizes $n_A$ and $n_B$. Let $t_A$ and $t_B$ denote the number of generations of vegetative growth from a single founder cell for populations $A$ and $B$, respectively. Their relative difference in growth initiation, expressed in generation can be written as

$$\Delta t_{A,B} = t_A - t_B = \log_2(n_A) - \log_2(n_B) = \log_2\left(\frac{n_A}{n_B}\right).$$

Thus, under the assumption of identical growth rates, the ratio of the sizes of two populations provides a direct measure of the difference in their growth initiation times.

If populations $A$ and $B$ are part of a larger population of size $n_{\text{tot}}$, their relative frequencies are

$$p_A = \frac{n_A}{n_{\text{tot}}} \quad \text{and} \quad p_B = \frac{n_B}{n_{\text{tot}}}.$$

We therefore have

$$\log_2(p_A) - \log_2(p_B) = \log_2\left(\frac{p_A}{p_B}\right) = \log_2\left(\frac{n_A/n_{\text{tot}}}{n_B/n_{\text{tot}}}\right) = \log_2\left(\frac{n_A}{n_B}\right) = \Delta t_{A,B}.$$

Importantly, the total population size cancels from the ratio, allowing the relative difference in growth initiation to be estimated directly from the measured population frequencies.

#### Delay computation in the $Z = 1$ tetrads

We next apply this framework to the population produced from $Z = 1$ tetrads. Consider, for example, the [G-R] and [G-G] population - the same reasoning applies to the other pairwise comparison. We can define the corresponding delay as

$$\Delta t_{[G-R],[G-G]} = \log_2(p_{[G-R]}) - \log_2(p_{[G-G]})$$

In an idealized scenario in which the [G-R] and [G-G] populations each originate from a single zygote, $\Delta t_{[G-R],[G-G]}$ represents the difference, expressed in mitotic generations, between the initiation of vegetative expansion of the intra-tetrad mating lineage and the one of the green-spore haplo-selfing lineage.

This is the quantity reported in the main results of the paper, and provides a simple and interpretable way to express differences in the final contribution of the two mating strategies as equivalent difference in the timing of vegetative growth initiation.

However, the early events following spore germination are likely more complex than represented by this idealized scenario. In particular, the green-spore haplo-selfing lineage may not originate from a single zygote. Therefore, rather than representing a literal difference in the timing of two mating events, $\Delta t_{[G-R],[G-G]}$ , is interpreted as an "effective" difference in lineage-expansion timing, expressing the observed difference in the contributions of the two mating strategies as an equivalent difference in the number of generations ("delay") of vegetative growth.

#### Interpretations of the delay under different scenarios

What values should we expect for $\Delta t_{[G-R],[G-G]}$?

Under the classical model of haplo-selfing (Haber, 1998), two successive mitotic cycles occur following spore germination before mating-type switching generates cells of opposite mating types that can subsequently mate. In our framework, we operationally define the onset of haplo-selfing lineage expansion as the first budding event, rather than the subsequent mating-type switching and cell-fusion events. Assuming synchronous germination, identical mitotic growth rates, and negligible duration of the mating-type switching process compared to the mitotic cycle, the intra-tetrad lineage combines the contribution of two spores, whereas each haplo-selfing lineage derives from a single spore. This two-fold difference corresponds to one mitotic generation, giving the baseline expectation for the relative expansion of the intra-tetrad and haplo-selfing lineages.

Thus, under the hypotheses described above, we expect

$$\Delta t_{[G-R],[G-G]} \simeq 1.$$

This expectation does not depend on the precise number of haploid divisions preceding mating. Even if additional vegetative growth occurs before mating, as observed for some germinating spores that enter the cell cycle rather than mate immediately (McClure et al., 2018), the initial two-fold difference in the contributions of the intra-tetrad and haplo-selfing lineages is preserved. Therefore, defining lineage expansion from the first budding event avoids assumptions about the precise sequence of events preceding mating. If zygote formation were instead used as the common starting point, the interpretation would differ. Under the classical haplo-selfing scenario, a single spore lineage can produce two zygotes, compared with one [G-R] zygote produced by intra-tetrad mating. This two-fold difference in founder number would correspond to an additional one-generation correction. The first-budding definition used here avoids this dependence on the number of zygotes subsequently contributing to each lineage.

Accordingly, ($\Delta t_{[G-R],[G-G]} \simeq 1$) represents the baseline expectation. Values above or below 1 indicate a greater or smaller effective expansion advantage of the [G-R] lineage, respectively.

Such deviations may reflect differences in germination timing or in the timing of subsequent events along the two mating pathways. We therefore interpret $\Delta t_{[G-R],[G-G]}$ as an effective difference in lineage-expansion timing, expressed in mitotic generations, rather than as a direct chronological delay between mating events.
